# Integrated nano-immunotherapy for synergizing cancer vaccination with sustained cytokine release and tumor microenvironment modulation

**DOI:** 10.64898/2026.11.26.769487

**Authors:** Zahara Thaivalappil Zakkariya, Saira Nujoom Mohammed, Margaux Saillard, Anjali Kozhissery Sunilkumar, Alejandra Gomez-Cadena, Hajar El Ahanidi, Sreedevi Radhakrishnan, Amal George, Amrita Murali Nair, Shantikumar Nair, Pedro Romero, Camilla Jandus, Manzoor Koyakutty

## Abstract

**Background:** Cancer immunotherapy has transformed clinical oncology, yet durable responses remain limited to approximately 25% of patients due to tumor heterogeneity and an immunosuppressive tumor microenvironment (TME). The barriers that underlie this limited efficacy include poor antigen presentation, inadequate cytokine signaling, and myeloid-derived suppressor cell (MDSC)-mediated immunosuppression. We hypothesized that a rationally engineered, tri-modal nano-immunotherapy addressing all three barriers simultaneously would produce more robust and durable antitumor immunity than single-agent approaches.

**Methods:** We developed a three-component nanotech platform: (i) nITC1, a tri-metal (Ca²⁺/Mn²⁺/Zn²⁺) hydrogen phosphate (n-CMZ) nanoparticle adjuvant-based vaccine synthesized by co-precipitation and loaded with tumor antigens; (ii) nITC2, a polyvinyl alcohol (PVA) based hydrogel enabling sustained release of the cytokine IL-15; and (iii) nITC3, a hydrogel delivering low-dose 5-fluorouracil (5-FU) to deplete MDSCs. Components were characterized by DLS, SEM, XRD, and ICP-AES, and evaluated *in vitro* using dendritic cell (DC) activation, the stimulator of interferon genes (STING) pathway (qRT-PCR/ELISA), and T-cell proliferation/IFN-γ ELISpot assays. *In vivo* efficacy was assessed in C57BL/6 mice bearing B16F10 melanoma, including subcutaneous and metastatic models, with immune profiling by flow cytometry.

**Results:** Nanoparticle adjuvant activated the cGAS-STING pathway, increasing IFN-β and CD80/CD86 expression in DCs, enhanced IFN-γ secretion from healthy human peripheral blood mononuclear cells (PBMCs), and promoted antigen-specific CD8⁺ T-cell proliferation. The nCMZ nanoparticle drove robust DC activation and antigen presentation. nITC2 sustained IL-15 release and expanded CD8⁺ T and NK cell populations, while nITC3 reduced MDSC frequencies, lowered TGF-β, and promoted M1 macrophage polarization. The combined nITC(1+2+3) regimen produced the greatest tumor growth inhibition, prolonged survival, and most effectively suppressed pulmonary metastases relative to monotherapy or dual-therapy controls.

**Conclusions:** Synchronized delivery of a nano-adjuvant vaccine, sustained cytokine support, and MDSC-targeted TME remodelling produced durable antitumor immunity exceeding that of single-modality vaccination. This tri-modal nanoplatform offers a translatable blueprint for combination cancer immunotherapy, warranting further evaluation of safety, biodistribution, and synergy with checkpoint blockade.

- **What is already know on this topic:** Cancer immunotherapy benefits only ∼25% of patients due to poor antigen presentation, weak cytokine support, and MDSC-driven immunosuppression. Systemic toxicity and short drug half-lives further limit conventional single-agent therapies.
- **What this study adds:** This study establishes a tri-modal platform integrating a tri-metal nano-vaccine, a sustained IL-15 hydrogel, and an MDSC-depleting 5-FU hydrogel. Together, they simultaneously activate dendritic cells, expand CD8⁺ T/NK cells, and remodel the tumor microenvironment.
- **How this study might affect research, practice or policy:** It provides a translatable framework for multi-target nano-immunotherapies that significantly improve tumor clearance and metastasis control. It also provides a rationale for testing synchronized delivery systems alongside checkpoint inhibitors in future clinical trials.

## Introduction

Cancer immunotherapy, which harnesses the immune system’s ability to combat cancer has seen remarkable advancements in the last decade^1^. Clinical evidences suggests that current immuno-therapeutics are effective in ∼ 25% of patients, leaving a substantial proportion with little or suboptimal responses^1–3^. Multiple factors contribute to this challenge, including tumor-immune heterogeneity and a highly immunosuppressive TME, which remain major barriers to durable antitumor immunity^4^. The TME has traditionally been viewed as a localized niche. However, growing evidence suggests that systemic immune dysfunction initiated early during tumorigenesis and sustained during metastasis is a key driver of tumor associated immune compromise^5^. Cancer is a systemic disease, with immune failure established not only in the TME but also in distant organs including bone marrow, spleen, lymph nodes, and the circulatory system^6^. Recent meta-analyses involving over 40,000 patients demonstrate that elevated circulating MDSCs and neutrophils are strongly associated with poor prognosis^7–9^. The preferential dissemination of tumors through lymph nodes, despite their central role in initiating anti-tumor immune responses, underscores the extent of systemic immune dysfunction in cancer^10^. While current immunotherapies predominantly focus on T-cell activation, they largely overlook the pivotal roles of MDSCs and M2 macrophages, which sustain immunosuppression within both the TME and peripheral immune system^11,12^.

A key challenge limiting cancer immunotherapy is the poor immunogenicity of tumor-associated antigens and their inadequate presentation by the innate immune system^13,3^. Most tumor antigens are derived from overexpressed self-proteins, leading to immune tolerance. Insufficient APC co-stimulatory signals (CD80/CD86/CD40, MHC-II) further limit effective immune activation^14^. Even when neo-antigens are presented, inferior co-stimulation may prevent effective T-cell activation^15^. Tumor-induced dysfunction of DCs further impairs immune response by disrupting their migration to lymph nodes and antigen processing and presentation^16,17^. Further, elevated levels of anti-inflammatory cytokines such as TGF-β and IL-10 in the TME and the lack of systemic pro-inflammatory cytokines such as IL-12, IL-15, IL-2 or IFN-γ also lead to weak immune response^18^.

In this study, we address these critical challenges of cancer immunotherapy using unique nanoengineered platforms. Although many nanomaterial systems, such as proteins, polymers ^19^, and lipids, are used in cancer drug delivery and mRNA vaccine delivery, they are least explored in cancer immunotherapy^20^. We developed a treatment platform comprising three complementary nano-immunotherapy components (n-ITCs): (i) a calcium-manganese-zinc phosphate nano-adjuvant cancer vaccine (nITC1) to enhance antigen delivery and innate immune activation, (ii) a PVA hydrogel for sustained IL-15 release to stimulate T and NK cells (nITC2), and (iii) the same hydrogel for sustained delivery of 5-FU (nITC3) to deplete MDSCs and M2 macrophages. In murine melanoma models, the integrated nITC platform enhanced antigen presentation, T-cell priming, and CD8⁺ T-cell infiltration while reducing local and systemic immunosuppression, leading to robust tumor regression. In metastatic melanoma, combined nITC treatment suppressed lung metastasis through enhanced CD8⁺ T-cell immunity and depletion of suppressive myeloid cells. Overall, the platform synergistically promoted anti-tumor immunity by enhancing antigen presentation, cytotoxic lymphocyte function, and remodelling the immunosuppressive TME.

## Materials and Methods

### Reagents

Calcium chloride (CaCl₂), manganese nitrate (Mn(NO₃)₂·xH₂O), zinc nitrate (Zn(NO₃)₂·xH₂O), diammonium hydrogen phosphate ((NH₄)₂HPO₄), trisodium citrate, ammonium hydroxide, and PVA were purchased from Sigma-Aldrich. Murine JAWS II DCs, RAW264.7 macrophages, and B16F10 melanoma cells were obtained from ATCC. PBMCs were isolated from buffy coats obtained from healthy donors at Amrita Institute of Medical Sciences, India. Dulbecco’s Modified Eagle Medium (DMEM), Roswell Park Memorial Institute medium 1640 (RPMI-1640), α-Alpha Minimum Essential Medium (α-MEM), fetal bovine serum (FBS) and penicillin-streptomycin were purchased from Gibco. Chemotherapeutic agents including 5-FU, gemcitabine, paclitaxel, cisplatin, and doxorubicin were purchased from Sigma-Aldrich.

### Synthesis and characterization of n-CMZ nanoparticles

Manganese (n-M), calcium (n-C), and zinc (n-Z) hydrogen phosphate nanoparticles were synthesized by wet-chemical precipitation method as described in our previously reported work ^21,22^. Optimized n-CMZ nanoparticles were prepared by co-precipitation of calcium, manganese, and zinc at a 1:4:1 molar ratio. Particle size, polydispersity index, and zeta potential were determined by dynamic light scattering (Zetasizer Nano ZS, Malvern). Morphology was examined by scanning electron microscopy (SEM; JEOL), elemental composition was confirmed by SEM-energy dispersive X-ray spectroscopy (SEM-EDX). Crystallinity was analyzed by X-ray diffraction (XRD; PANalytical X’Pert Pro), and metal ion content was quantified by inductively coupled plasma atomic emission spectroscopy (ICP-AES). All characterization experiments were performed in triplicate.

### Synthesis of nITC2 and nITC3 hydrogels

Hydrogels were prepared using 13% (w/v) PVA dissolved in sterile, endotoxin-free water at 80-90 °C under continuous stirring (500 rpm) until a clear viscous solution formed. For nITC2, 1 µg recombinant IL-15(R&D Systems) was added and mixed thoroughly. For nITC3, 5-FU was incorporated at a dose equivalent to mice 100 mg/kg. Both formulations were homogenized and subjected to one freeze-thaw cycle for physical crosslinking (−80 °C overnight, followed by thawing at room temperature).

### Cell viability assay

Cytotoxicity of nITC1 and nITC3 formulations was evaluated by MTT assay in JAWS II cells and human PBMCs (nITC1) and RAW 264.7 macrophages (nITC3). After 24 h treatment, cell viability was determined by MTT reduction assay.

### Cell culture and immune cell isolation

Murine JAWS II DCs, RAW264.7 macrophages, and B16F10 melanoma were maintained under standard conditions (37°C, 5% CO₂). RAW264.7 and B16F10 cells were cultured in DMEM supplemented with 10% FBS and 1% penicillin-streptomycin. JAWS II cells were cultured in complete α-MEM medium with recombinant murine (rm)GM-CSF(GenScript). Bone marrow derived dendritic cells (BMDCs) were generated from femurs and tibias of C57BL/6 mice. BMDCs were cultured in complete RPMI-1640 medium and rmGM-CSF for seven days and harvested on day 7 for downstream assays. Splenocytes were isolated by mechanical dissociation of spleens through 70-μm strainers, followed by erythrocyte lysis and washing. Healthy human PBMCs were isolated by density-gradient centrifugation using Histopaque-1077 (Sigma) and cultured in complete RPMI-1640 medium.

### Flow cytometry

Tumors, spleens, lymph nodes, and blood samples were processed into single-cell suspensions. Tumor tissues were enzymatically digested with 50U/mL collagenase type I (Gibco) and 100µg/mL DNase I (STEMCELL Technologies), filtered through 70 µm strainers, and subjected to red blood cell lysis. Cells were stained with viability dye and fluorochrome-conjugated antibodies, and analyzed using a BD FACSLyric™ flow cytometer. Antibodies (CD45-APCR700, CD3-FITC, CD4-APC, CD8-PercpCy5.5, CD11b-FITC, CD11c-BV421, MНСII-РЕ, B220-BV605, Ly6G-BV711, Ly6C BV605, CD86-PE Cy7, CD206-AF647, CD49b-BV510, NK1.1-BV711, CD44-BV510, CD62L-BV786, CD69-PE-Cy7, CD40-BV510, CD80-BV786, F4/80-Percp Cy5.5, PD1-BV711, Fixable Viability stain 450) were purchased from BD Biosciences and BioLegend. Intracellular staining was performed using BD Cytofix/Cytoperm reagents. Data analysis was performed using FlowJo software.

### Nanoparticle uptake study

Fluorescent n-CMZ nanoparticles were prepared by incorporating Nuclear Fast Red (Abcam) during synthesis and subsequently conjugated with FITC-labeled ovalbumin (OVA-FITC, Invitrogen). JAWS II cells were incubated with fluorescent nanoparticles for 24 h before analysis. Cellular uptake was visualized by fluorescence microscopy (Nikon 80i) and quantified by flow cytometry.

### DC activation and antigen presentation assays

JAWS II cells and BMDCs were treated with nanoparticles or indicated formulations for 24 h. DC activation was evaluated by flow cytometric analysis of CD40, CD80, CD86, and MHC-II expression. For antigen presentation studies, BMDCs were incubated with OVA (5 µg/mL) following nanoparticle treatment, and OVA257-264-H-2Kb complex formation was assessed using PE-conjugated 25-D1.16 antibody (Invitrogen). STING pathway activation was evaluated by qRT-PCR analysis of STING, IFIT1, IFIT2, and IFIT3 expression following RNA isolation (QIAGEN RNeasy Kit) and cDNA synthesis (iScript, Bio-Rad). Primer sequences used were: STING forward 5’-GCTCCAGGATGAACCTGATGA-3’, reverse 5’-TGATGCCAAGGAAGTTAGGGA-3’; IFIT1 forward 5’-CTCTGAAAGTGGAGCCAG AAAAC-3’, reverse 5’-AAATCTTGGCGATAGGCTACG A-3’; IFIT2 forward 5’-CTGAAGCTTGACGCGGTACA-3’, reverse 5’-ACTTGGGTCTTTCTTTAAGGCTTCT-3’; and IFIT3 forward 5’-TTCCCAGCAGCACAG AAA C-3’, reverse 5’-AAATTCCAGGTGAAATGGCA-3’. Relative gene expression was normalized to β-actin using the 2^-ΔΔCt method, and IFN-β secretion was quantified by ELISA (R&D Systems).

### T-cell activation and antigen-specific proliferation assays

Mouse splenocytes were activated with anti-CD3 and anti-CD28 for 24 h and treated with nanoparticles. Cells were stained for CD4/CD8/CD40L expression and analyzed by flow cytometry. For antigen-specific proliferation studies, BMDCs were treated with n-CMZ nanoparticles loaded with N4 peptide (12.5-50 µg/mL) for 24 h. CD8⁺ T-cells isolated from OT-1 mice were labeled with carboxyfluorescein succinimidyl ester (CFSE, Invitrogen) and co-cultured with treated BMDCs at a 1:5 BMDC ratio for 72 h. T-cell proliferation was quantified by CFSE dilution using flow cytometry.

### IFN-γ response study

Human PBMCs were treated with nanoparticles for 24 h. IFN-γ-producing cells were quantified using a human IFN-γ ELISpot kit (Mabtech). For ex vivo immune response analysis, splenocytes from treated C57BL/6 mice were restimulated with 5 µg gp100/TRP2 peptide (GenScript) and analyzed using a mouse IFN-γ ELISpot assay.

### nITC2 and nITC3 release kinetics

For in vitro kinetics of nITC2, hydrogels were prepared 13% and 15% w/v) loaded with BSA (500 µg) were evaluated for in vitro release in PBS (pH 7.4) over 48 h. Protein release was quantified at predefined time points by UV-Vis absorbance at 280 nm using a BSA standard curve. For in vivo kinetics, nITC2 hydrogels containing recombinant mouse IL-15 (1 µg) were administered subcutaneously to C57BL/6J mice, and plasma IL-15 levels were measured at 4, 28, 72, and 120 h using a Mouse IL-15 DuoSet ELISA kit (R&D Systems). nITC3 hydrogels containing 5-FU (100 mg/kg) were administered subcutaneously, and plasma 5-FU concentrations were quantified up to 168 h by HPLC using a C18 reverse-phase column with UV detection at 263 nm.

### Tumor model

Wild-type mice C57BL/6J were purchased from Charles Rivers, France for the experiments done in Switzerland, whereas those used in India were bred and housed at the Central Laboratory Animal Facility of Amrita School of Medicine, India. C57BL/6J mice (6–14 weeks old) were used for all studies. Mice were maintained in accredited animal facilities and all animal experiments were performed in compliance with relevant ethical guidelines for the use and the care of laboratory animals. A 12 h dark/light cycle, at 21 °C ± 1 °C and 55 ± 10% of relative humidity was ensured in the facilities. For the subcutaneous melanoma model, mice were inoculated with 1 × 10⁵ B16F10 cells in the right flank, and tumor volume was calculated as π/6 × length × width². For the metastatic model, mice received 5 × 10⁵ B16F10 cells intravenously and were treated according to the optimized schedule. At the study endpoint, lungs were collected for metastatic nodule assessment and immune profiling. Animal experiments were approved either by the Institutional Animal Ethics Committee of Amrita Institute of Medical Sciences, Kochi, India or by the Veterinary Authorities of the Swiss Geneva and Vaud Cantons under the authorizations VD3496 and VD3772.

### *In vivo* nITC2 activation

Healthy C57BL/6 mice received IL-15-loaded nITC2 hydrogel (1 µg IL-15), free IL-15, or vehicle control by subcutaneous injection. On days 2 and 7, blood, spleen, and draining lymph nodes were collected for immune profiling by flow cytometry.

### *In vivo* anti-tumor study and survival analysis

For nITC1 vaccine formulation optimization, tumor-bearing mice received TRP2 and gp100 (50 µg each) loaded nanoparticles containing different concentrations of Mn, Zn, and Ca. Optimized nITC1-CMZ nanoparticles contained calcium, manganese, and zinc at a 1:4:1 ratio and were administered subcutaneously. For nITC3 evaluation, tumor-bearing mice received free 5-FU or nITC3 hydrogel containing 5-FU (100 mg/kg). Tumors, immune organs and blood plasma were collected at endpoint for cytokine analysis and immune profiling. For combination therapy studies, B16F10 tumor-bearing mice received sequential treatment with nITC1, nITC2, and nITC3. Treatment was administered post tumor induction as follows: nITC1 on day 7, nITC2 on day 8, booster doses of nITC1 and nITC2 on days 12 and 13, and nITC3 on day 18. Control groups included untreated mice, individual components, dual therapy, and standard vaccine controls. A single-dose combination therapy was performed in the same tumor model. Mice received a sequential single-dose regimen of nITC1 (day 7), nITC2 (day 8), and nITC3 (day 13). Tumor progression and survival were monitored according to predefined humane endpoints.

### Western blot analysis

Tumor tissues were lysed in extraction buffer containing PMSF and protease inhibitor cocktail. Equal protein amounts were separated by SDS-PAGE, transferred onto PVDF membranes, and blocked with 5% non-fat milk. Membranes were incubated with IDO(D8W5E) Rabbit mAb and Arginase-1(D4E3M) Rabbit mAb (Cell Signaling Technology), followed by HRP-conjugated secondary antibody. Protein bands were detected using Bio-Rad Gel Doc system.

### Statistical Analysis

Data are presented as mean ± SEM. Statistical analyses were performed using GraphPad Prism 9 (GraphPad Software). Tumor growth was analyzed by one-way ANOVA with Tukey’s multiple-comparison test, while immune cell infiltration was analyzed using Dunnett’s multiple-comparison test and an unpaired two-tailed Student’s *t*-test. A *p* value < 0.05 was considered statistically significant.

## Results

### Engineering Ca-Mn-Zn-Phosphate nanoparticles as a multi-functional vaccine adjuvant

We developed n-CMZ designed to simultaneously activate the STING/IFN-β pathway and enhance antigen presentation and maturation in DCs. Building on our previous work using doped calcium phosphate nanoparticles for cancer-imaging and drug delivery ^21,22^, here, we engineered an immunologically functional nano-adjuvant by co-mineralizing Ca²⁺, Zn²⁺, and Mn²⁺ ions, at optimized doses, suitable for activating DCs and T-cells (Figure 1A). SEM images of particles showed an irregular morphology with a size distribution between 100-1000 nm (Figure 1B) with an average hydrodynamic diameter of 260.1 nm (Figure 1C) and a zeta potential of −12.1 mV (Supplementary Figure 1A). XRD analysis indicated a largely amorphous structure (Figure 1D), while ICP analysis quantified the elemental concentration of metal ions (Figure 1E) and confirmed by EDX analysis (Supplementary Figure 1B). The optimized nano-adjuvant, n-CMZ, comprises calcium (n-C), manganese (n-M), and zinc (n-Z) nanoparticles. To assess cellular delivery, DCs were co-cultured with fluorescently labelled n-CMZ (red) complexed with OVA-FITC. Confocal microscopy showed efficient uptake with intracellular co-localization of antigen and adjuvant (yellow fluorescence) in the cytosol (Figure 1F) as also confirmed by flow cytometry (Supplementary Figure 1C). Cytotoxicity of n-C, n-M, and n-Z was evaluated in mouse DCs (Figure 1G) and human PBMCs (Supplementary Figure 1D) across 10-4000 µM. MTT assays showed dose-dependent effects, with high viability at lower doses and ∼60-70% viability only at 2000-4000 µM. All formulations showed IC₅₀ > 4000 µM in both cell types. Given the established role of Mn in activating the cGAS-STING pathway ^23–25^, nCMZ was optimized with varying Mn concentrations to STING agonistic activity. Treatment of DCs with individual n-M, n-Z, or n-C nanoparticles revealed that n-M (100 μM Mn) strongly upregulated STING expression in JAWS II cells, surpassing the response induced by cGAMP (Figure 1H). n-M also induced significantly higher IFN-β secretion than n-C and n-Z (Figure 1I). This was further corroborated by increased mRNA expression of STING-regulated genes IFIT-1, IFIT-2, and IFIT-1^26^ in BMDCs treated with n-M (Supplementary Figure 1E). n-M also promoted DC maturation, as evidenced by greater CD86 expression than that induced by LPS (Supplementary Figure 1F). We then examined the immunomodulatory contributions of Zn and Ca, which are extensively investigated micronutrients owing to their pivotal roles in immune function and protein biochemistry^27–29^. Given the established role of Zn in T-cell function^30^, IFN-γ secretion was assessed in PBMCs following treatment with n-Z, n-C, and n-M. At 100 μM, n-Z induced significantly higher IFN-γ secretion than n-C or n-M (Figure 1J). To evaluate the effects of these elements on antigen presentation and T-cell priming, BMDCs pulsed with N4-OVA were treated with these nanoparticles (12.5-50 μM) and co-cultured with OVA-specific OT-I T-cells. OT-I proliferation increased with n-C treatment, whereas n-Z and n-M showed no comparable effect (Figure 1K). n-M treatment (25-125 μM) also enhanced MHC-II, CD86, and CD40 expression in DCs (Figure 1L). In contrast, n-C induced dose-dependent increases in MHC-II and CD40 without affecting CD86, indicating a mechanistically distinct role for Ca²⁺. n-Z produced minimal changes in CD86 and modest increases in MHC-II and CD40, suggesting a more selective immunomodulatory effect for each element. Next, mouse splenocytes were co-cultured with nanoparticles for 96 h and T-cell activation markers were quantified. In CD8 T-cells, both n-M and n-Z significantly increased CD40L expression in a dose-dependent manner between 12.5 and 25 μM, whereas n-C induced only modest, dose-independent effects (Figure 1M). In CD4 T-cells, all particles produced dose-dependent increases in CD40L⁺ populations (Figure 1N). These findings demonstrate element-specific and dose-dependent modulation of DC and T-cell activation by n-CMZ nanoparticles, highlighting the importance of optimizing Ca, Zn, and Mn composition for synergistic immune activation.

**Figure 1.**
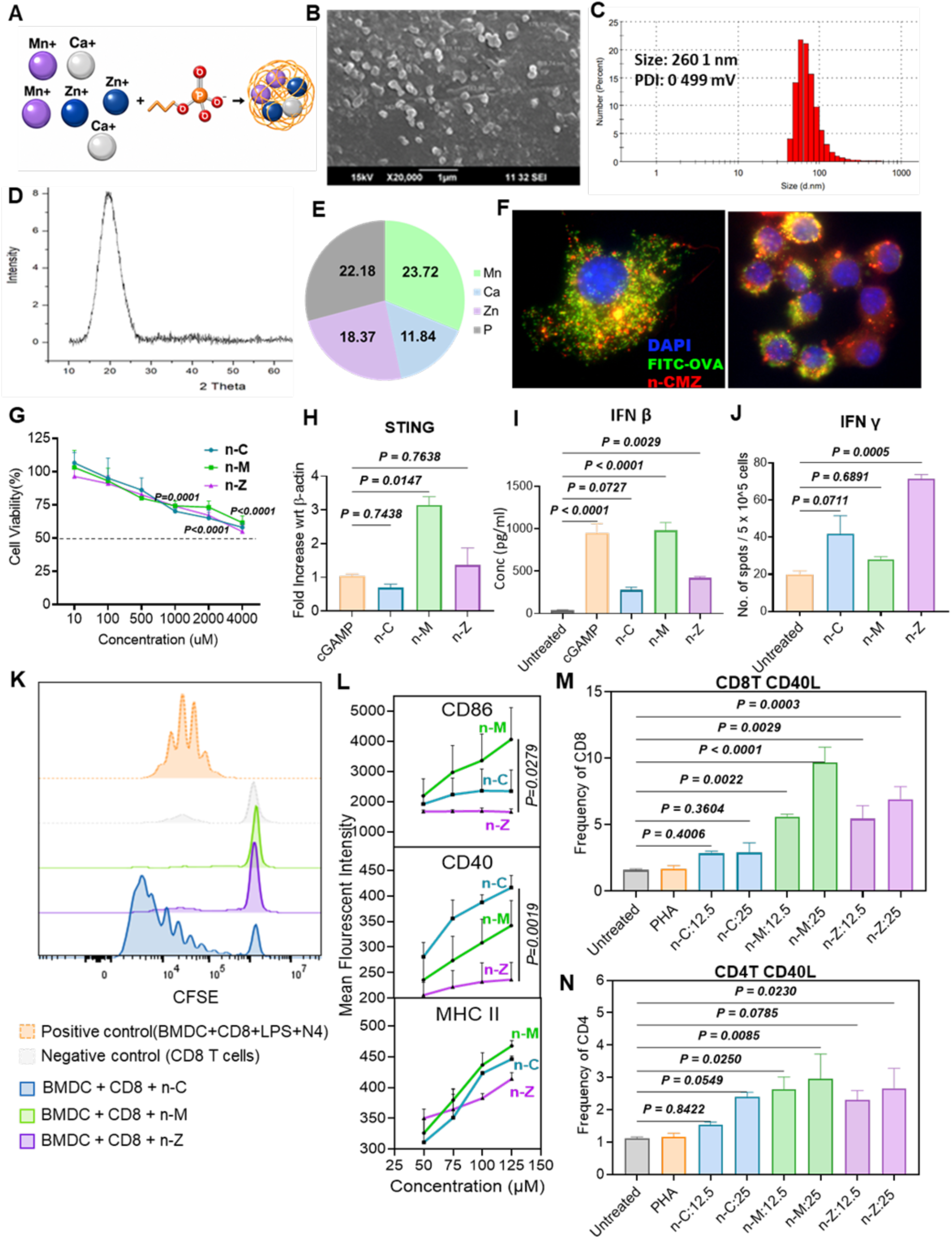
Optimized composition of nano-adjuvant (n-CMZ) modulates DC and T cell activation. A) Schematic illustration showing the formation of a combination of Ca, Mn, Zn containing tri-metal phosphate adjuvant nanoparticles (n-CMZ). B) SEM images of n-CMZ particles showing size distribution. C) Dynamic light scattering-based hydrodynamic size measurement of n-CMZ particles. D) X-Ray diffraction pattern of n-CMZ particles. E) ICP-MS analysis data showing the weight fraction (mg/g) of Ca, Mn, Zn in n-CMZ. F) Fluorescent microscopic images showing DC uptake of nanoparticles, stained in red, conjugated with OVA-FITC peptide (green) at 60X (left) and 40X(right) magnification. G) Dose-dependent cytotoxicity of nanoparticles containing individual elements in the mouse DC line, JAWS II. H) *STING* mRNA expression in the JAWS II cell line treated with n-Z, n-M, and n-C nanoparticles. I) *IFN-β* expression studies in JAWS II cell line treated with n-Z, n-M, and n-C nanoparticles. J) IFN-γ ELISPOT in human PBMCs treated with n-Z, n-M, and n-C nanoparticles. K) CFSE assay showing proliferation profile of OT-1 cells co-cultured with different nanoparticles-treated BMDCs. L) DC response of CD40, CD86, MHC-II with different concentrations of n-C, n-M, and n-Z nanoparticles. M-N) Bar graph showing dose-response of CD40L expression in CD8 and CD4 T cells. Data are represented as mean ± SEM. P values in (G, L) were generated using two-way ANOVA with Tukey’s multiple comparison test. P values in (H, I, J) and (M, N) were generated using ordinary one-way ANOVA with Tukey’s multiple comparison test for comparison. Panel A was created using BioRender.

### Ion-driven modulation of DC activation and anti-tumor immunity via engineered nanoparticles

Following *in vitro* characterization, the optimized nano-adjuvants were evaluated in murine melanoma vaccination studies. Vaccine formulations containing single-ion (n-M, n-Z, n-C), bimetallic (n-MZ), or trimetallic (n-CMZ) nano-adjuvants with TRP2 and gp100 antigens (nITC1-M, nITC1-MZ, and nITC1-CMZ) were administered on days 7, 12, and 17 after tumor implantation (Figure 2A). The immunostimulatory role of n-M was first evaluated by administering 1-, 2-, or 4-mM Mn with fixed doses of TRP2 and gp100. n-M elicited a dose-dependent anti-tumor response, with optimal tumor inhibition and tumor-infiltrating lymphocyte infiltration observed at 2 mM (Figure 2B; Supplementary Figure 2A). Next, immunological synergy between Mn and Zn ions were examined. We hypothesize that Zn could boost vaccine effectiveness by activating IFN-γ signaling in T-cells ^31^, while Mn enhances innate immune responses in myeloid cells^32^. Bimetallic nanoparticles containing 2 mM Mn and varying Zn concentrations (0.5, 1, or 2 mM) were complexed with TRP2 and gp100 to generate nITC1-MZ vaccines. The Mn:Zn ratio of 4:1 (2 mM Mn, 0.5 mM Zn) induced the strongest antitumor response (Figure 2C), with increased infiltration of CD8^+^ T-cells, CD4^+^ T-cells, and NK cells in both tumor and spleen (Supplementary figure 2(I)B,C). Higher Zn doses (1 and 2 mM) reduced efficacy, likely due to Zn-induced immune stress and reduced cell viability^33^. These *in vivo* results were supported by *in vitro* assays using BMDCs, where a n-M: n-Z ratio of 4:1 induced the highest expression of activation markers CD86, CD40, CD80, and MHC-II in a dose-dependent manner (Supplementary Figure 2(II)A). Ca²⁺ concentration was titrated within the optimized n-MZ (4:1) framework to generate the final tri-metallic n-CMZ nanoparticles. In BMDCs, moderate Ca²⁺ levels enhanced DC activation, whereas higher doses reduced CD40 and CD86 expression. The optimal immunostimulatory composition was identified as Ca:Mn:Zn = 1:4:1, using standardized doses of n-M (100 µM) and n-Z (25 µM), with n-C ranging from 25-100 µM (supplementary figure 2(II)B). Cytotoxicity of n-CMZ (1:4:1) was assessed in mouse DCs and PBMCs by MTT assay (Supplementary Figure 2(III)A). Across increasing elemental concentrations (Ca 12.5-500 µM, Mn 50-2000 µM, Zn 12.5-500 µM), cell viability decreased in a dose-dependent manner, reaching ∼60% at the highest combined dose (Ca 500, Mn 2000, Zn 500 µM). No IC₅₀ was reached, and the highest non-cytotoxic dose (500:2000:500 µM) was selected for *in vivo* studies. An OVA antigen presentation assay further showed enhanced antigen presentation by both CD11c⁺ and CD11b⁺ BMDCs (Supplementary Figure 2(III)B,C). In a B16F10 melanoma model, n-CMZ (1:4:1; Mn 2 mM, Zn 0.5 mM, Ca 0.5 mM) significantly reduced tumor growth compared to controls and n-MZ (Figure 2D). Flow cytometry analysis revealed increased intratumoral CD8^+^ and CD4^+^ T-cells (Figure 2E), along with enrichment of effector memory subsets (Figure 2F-G) and elevated activation markers including KLRG1 and CD69 (Figure 2H-K). These findings demonstrate that precise tuning of Mn, Zn, and Ca within the phosphate nanoparticle system enhances both innate and adaptive immune responses, with Ca²⁺ further boosting T-cell activation and antitumor efficacy.

**Figure 2.**
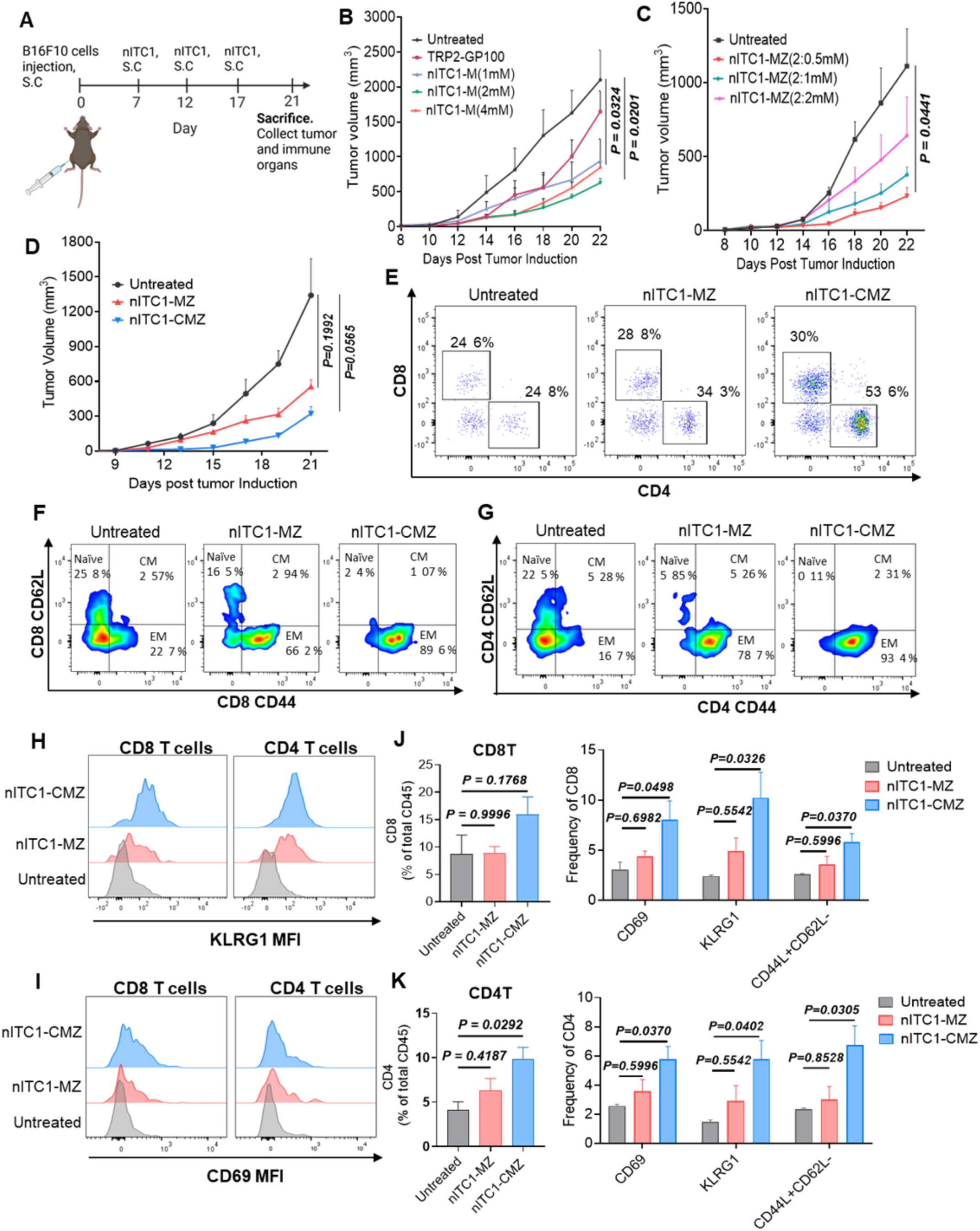
nITC1 *in vivo* and in vitro characterization. A) Schematic representation of *in vivo* melanoma tumor study in C57BL/6J. B) Tumor curve showing anti-tumor response of different concentration of n-M in nITC1; n=3; Experimental units=2. C) Tumor growth curves showing the anti-tumor response following incorporation of different concentrations of n-Z (0.5, 1, and 2 mM) into n-M (2 mM) nanoparticles to form n-MZ; n=4; Experimental units=2. D) Tumor growth curves comparing the anti-tumor efficacy of nITC1-MZ (4:1) and nITC1-CMZ (1:4:1) formulations in a melanoma tumor model, where n-M = 2 mM and n-Z = 0.5 mM for nITC1-MZ, and n-C = 0.5 mM, n-M = 2 mM, and n-Z = 0.5 mM for nITC1-CMZ; n=4; Experimental units=2. E) Representative flow cytometry plots showing CD4^+^ and CD8^+^ T cell populations in untreated, nITC1-MZ (4:1), and nITC1-CMZ (1:4:1) groups. F-G) Representative flow cytometry scatter plots showing central memory (CM) and effector memory (EM) subsets of CD8^+^ and CD4^+^ T cells in untreated, nITC1-MZ (4:1), and nITC1-CMZ (1:4:1) groups. H-I) Overlayed histograms showing KLRG1 (H) and CD69 (I) expression in CD8^+^ and CD4^+^ T cells in the nITC1-CMZ group compared to untreated and nITC1-MZ groups. J) Bar graphs showing frequency of CD8^+^ T cell infiltration in tumor and their activation across untreated, nITC1-MZ (4:1), and nITC1-CMZ (1:4:1) groups. K) Bar graphs showing frequency of CD4^+^ T cell infiltration in tumor and their activation across untreated, nITC1-MZ (4:1), and nITC1-CMZ (1:4:1) groups. Data are represented as mean ± SEM. P values in (B, C, D) were generated using ordinary one-way ANOVA with Dunnet’s multiple comparison’s test for comparison and (J, K) by ordinary one-way ANOVA with Tukey’s multiple comparison’s test for comparison. n = number of animals per group. Panel A was created using BioRender.

### Hydrogel-based nITC2 sustains release of IL-15 to augment immune response

Despite decades of research, only a limited number of cytokine therapies, such as IL-2 and IFN-α/β, have received FDA approval. Their clinical utility remains constrained by systemic toxicity and the inability of bolus administration to maintain sustained immune activation^34^. This highlights the need for sustained delivery systems that minimize systemic exposure while maintaining immune stimulation for 1-2 weeks after a single administration^35^. Here, we developed nITC2, an injectable sustained-release cytokine hydrogel. After comparing IL-2, GM-CSF, IL-12, and IL-15, IL-15 was selected due to its favourable safety profile and its ability to promote the survival, expansion, and cytolytic function of NK and CD8^+^ T-cells^36–38^. Importantly, unlike IL-2, IL-15 does not induce regulatory T-cells or cause vascular leak syndrome, a major dose-limiting toxicity associated with IL-2^39,40^. To generate the injectable nITC2 depot, we used FDA-approved biocompatible PVA^41^ using a scalable freeze-thaw cross-linking method (Figure 3A). SEM and micro-CT analysis revealed a porous hydrogel architecture with porosity inversely correlated with PVA concentration (8-13 wt%), enabling tuneable cytokine release kinetics (Figure 3B-C). For *in vivo* release studies, nITC2 was administered subcutaneously to C57BL/6J mice, and plasma IL-15 levels were measured at 4, 28, 72, and 120 h. IL-15 concentrations peaked at 4 h and gradually declined over 120 h (Figure 3D). Immunological assessment at days 2 and 7 (Figure 3E) demonstrated that nITC2 significantly increased peripheral CD4^+^ and CD8^+^ T-cell frequencies and upregulated activation markers (CD44, CD69, and PD-1) at day 2 compared with untreated and free IL-15 controls (Figure 3F-G). NK-cell frequency and activation were also enhanced in the nITC2 group (Figure 3H). By day 7, these responses returned to baseline, indicating a transient but controlled immune activation profile. In draining lymph nodes and spleen, nITC2 similarly increased NK-cell frequency and activation (Figure 3I), with transient CD8^+^ T-cell activation also observed (Supplementary figure 3(I)-3(II)). These findings demonstrate that nITC2-mediated IL-15 delivery induces sustained yet self-limiting systemic immune activation.

**Figure 3.**
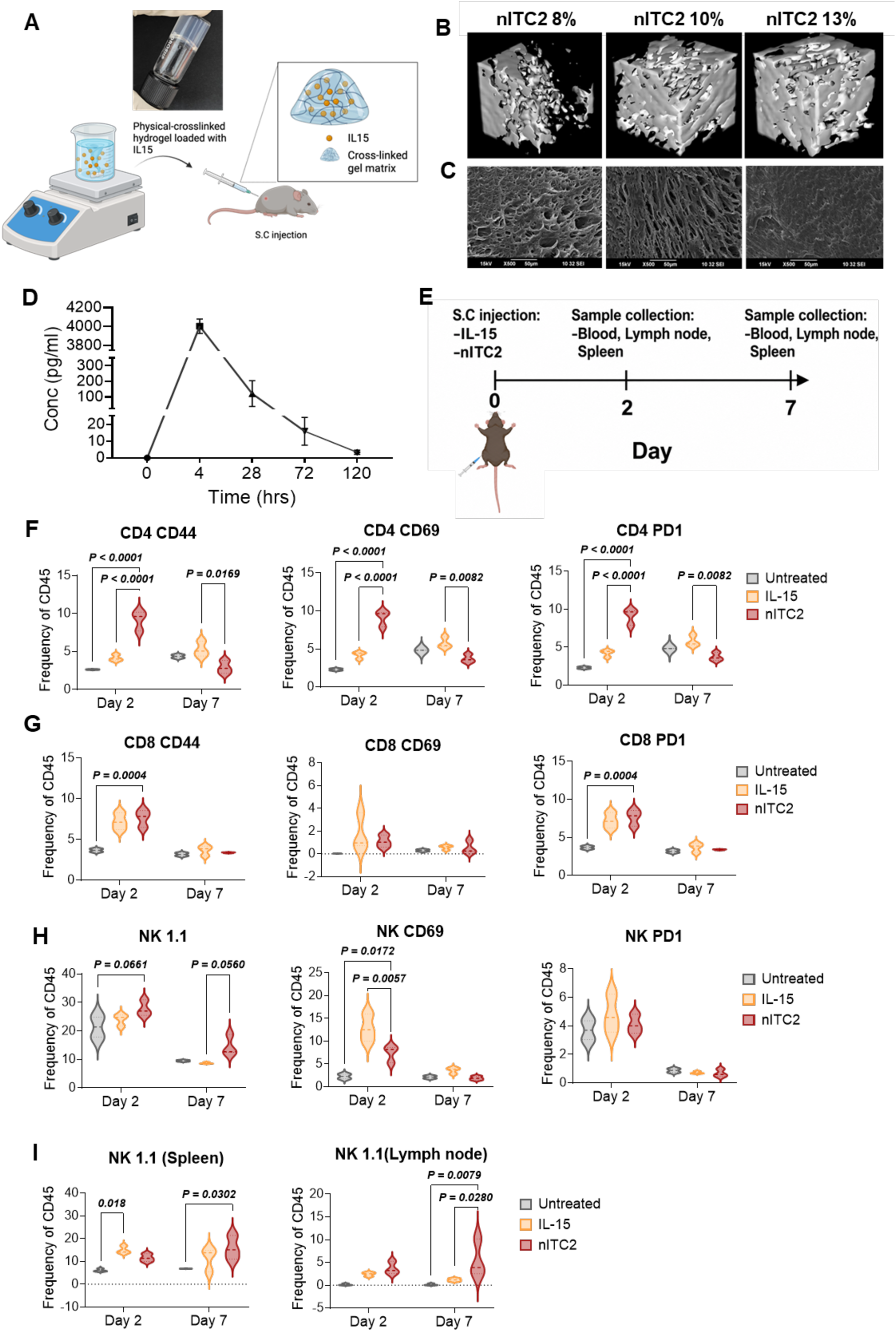
nITC2 preparation and characterization. A) Preparation of nITC2. B) Micro-CT image showing the porosity of nITC2 at different gel percentages. C) SEM image of nITC2 at different gel percentages. D) *In vivo* release of nITC2 loaded with IL-15 in C57 BL/6J mice; n=3; Experimental units=2 E) Schematic diagram showing treatment schedule of nITC2 activation study in healthy mice. F-H) Bar graphs showing the frequency of CD4^+^ T, CD8^+^ T and NK cell activation (CD44, CD69 and PD1) in peripheral blood on day 2 and day 7 post-treatment. I) Bar graphs showing the frequency of total NK cell populations in spleen (left) and lymph node (right); n=3; Experimental units=2. Data are represented as mean ± SEM. P values were generated using ordinary one-way ANOVA with Tukey’s multiple comparison’s test for comparison. p < 0.05. n = number of animals per group. Panels A and E were created using BioRender.

### nITC3 hydrogel targeting MDSCs to overcome immune suppression in the TME

MDSCs are a major obstacle for effective immunotherapy due to their ability to suppress cytotoxic T-cell activity, promote Tregs, and facilitate tumor progression^42,43^. We previously demonstrated that 5-FU can selectively deplete MDSCs in a PD-1-resistant Lewis lung carcinoma model^44^. We validated this finding in melanoma with other small molecules (Supplementary Figure 4A-C), where 5-FU showed the most substantial MDSC depletion and enhanced infiltration of CD4^+^ and CD8^+^ T-cells, compared to gemcitabine or chlodronate, which correlates with previous studies^45^. Clinically approved dose of 5-FU ranges between 400 to 3000mg/m^2^ (mouse equivalent: 133-1000mg/Kg)^46^. We used a lower dose of 100 mg/kg in the mouse studies, at which an anti-MDSC effect was observed^44^. Further, to overcome the challenge posed by the short half-life (4.5 to 13 mins) of 5-FU ^46^, we developed a sustained-release, injectable hydrogel using PVA, termed as nITC3 hereafter. Figure 4A illustrates the experimental design of nITC3, consisting of 13 wt% PVA loaded with 5-FU, administered subcutaneously in mice. SEM analysis (Figure 4B) revealed a porous, uniformly distributed microstructure in the freeze-dried hydrogel. *In vivo* pharmacokinetics showed that serum 5-FU peaked at 2 h post-injection and remained detectable up to 7 days, confirming sustained release (Figure 4C). To evaluate efficacy, nITC3 was tested in a B16F10 melanoma therapeutic model (Figure 4D). When tumors reached ∼100 mm³, mice received a single dose of either free 5-FU or nITC3 (100 mg/kg, 200 µL, s.c.). nITC3 produced significant tumor suppression compared to PBS and free drug controls (Figure 4E). Significant depletion of intratumoral PMN and M-MDSCs (Figure 4F-G), accompanied by macrophage repolarization toward an M1 phenotype (∼29%) and a reduction in M2 macrophages (∼4-fold decrease) (Figure 4F-H) were observed. This was associated with increased tumor-infiltrating CD8^+^ (48%) and CD4^+^ (41.5%) T-cells, compared to 17.5% and 2.73% in controls, respectively (Figure 4F-I). Activated CD4^+^ and CD8^+^ T-cells were also increased ∼2-fold (Figure 4J). nITC3 also reduced systemic MDSCs and promoted M1/M2 repolarization in the spleen (Supplementary Figure 4D). Intratumoral NK and B cells were also increased (Supplementary Figure 4E). In addition, TGF-β levels in tumor tissue and plasma were significantly reduced (Figure 4K), consistent with MDSC depletion. Western blot analysis further showed decreased IDO and Arg-1 expression in tumors (Supplementary Figure 4F). Overall, nITC3-mediated sustained 5-FU release (∼7 days) effectively depleted MDSCs systemically and locally, reversed immunosuppression, and enhanced CD4^+^ and CD8^+^ T-cell infiltration.

**Figure 4.**
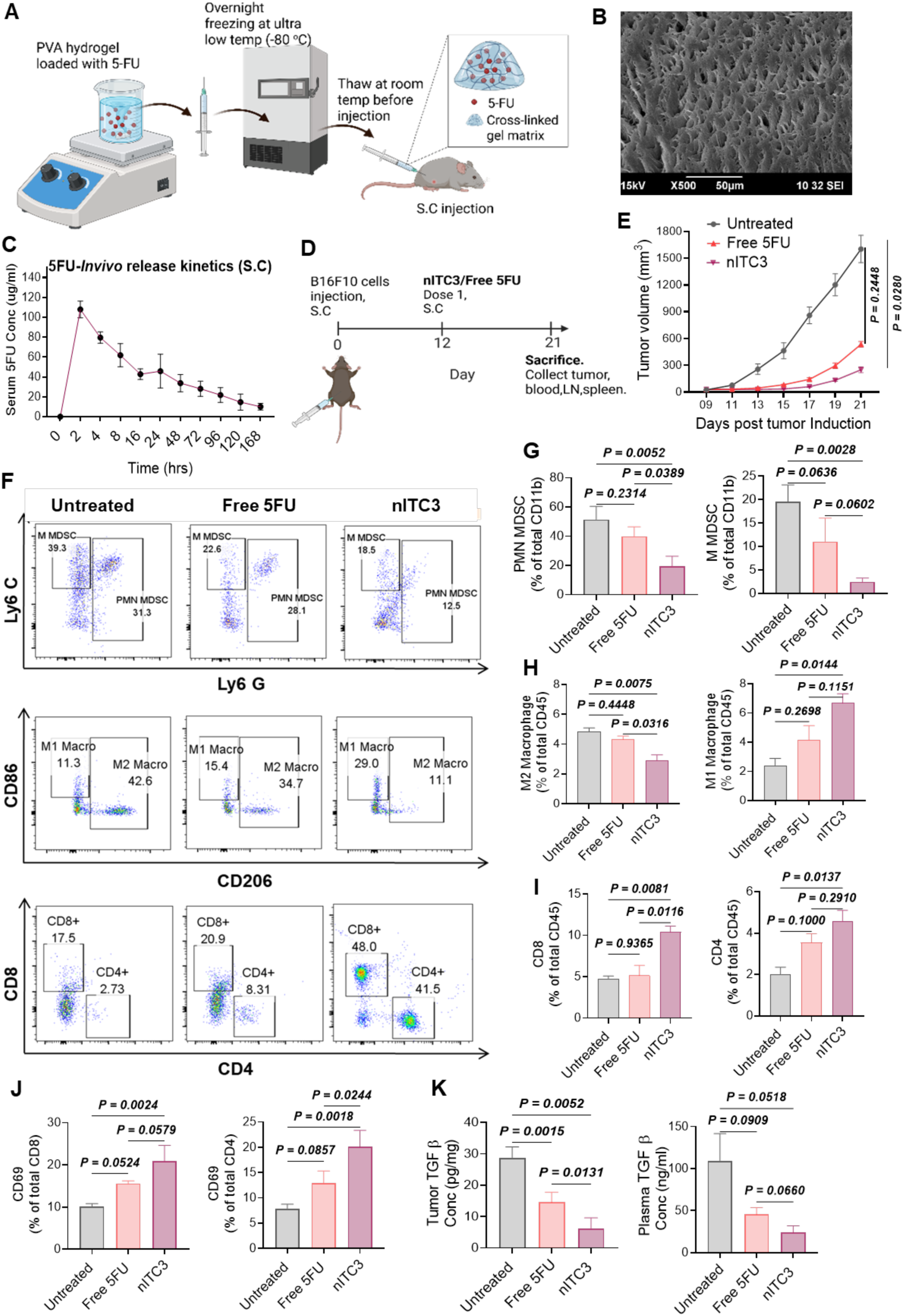
Immunomodulatory effects of nITC3 hydrogel in a melanoma tumor model. A) Schematic illustration of the preparation of the nITC3 hydrogel, encapsulation of 5-FU, and its subcutaneous administration into tumor-bearing mice. B) SEM image of nITC3. C) *In vivo* release kinetics of 5-FU from the nITC3 formulation over time n=3; Experimental units=2. D) Experimental timeline depicting melanoma tumor induction, treatment initiation, and analysis endpoints. E) Tumor growth curves comparing untreated, free 5-FU, and nITC3-treated groups; n=4; Experimental units=2. F) Representative flow cytometry plots showing M-MDSCs and PMN-MDSCs (top panel), tumor-associated macrophages (middle panel), CD4^+^ and CD8^+^ T cells in untreated controls, free 5-FU and nITC3 treated mice (bottom panel). G) Bar graphs showing PMN-MDSCs (left) and M-MDSCs (right) frequencies in peripheral blood in untreated controls, free 5-FU and nITC3-treated mice. H) Bar graphs showing M2 Macrophages (left) and M1 Macrophages (right) frequencies in peripheral blood in untreated controls, free 5-FU and nITC3-treated mice. I) Bar graphs showing intratumoral CD8^+^ (left) and CD4^+^ (right) T cells following nITC3 therapy relative to free 5-FU. J) Bar graphs showing frequency of intratumoral CD8^+^ CD69 (left) and CD4^+^ CD69 (right) T cells following nITC3 therapy relative to free 5-FU. K) TGF-β levels within the TME (left) and in the circulation (right) following nITC3 treatment, compared with free 5-FU. Data are represented as mean ± SEM. P values were generated using ordinary one-way ANOVA with Dunnet’s multiple comparison’s test for comparison. n = number of animals per group. Panel A and D were created using BioRender.

### Nano-vaccination with sustained cytokine delivery and TME myeloid modulation

We developed an integrated nano-immunotherapeutic platform combining optimized nITC1, nITC2, and nITC3 components to evaluate synergistic antitumor activity. In C57BL/6 mice bearing subcutaneous B16F10 melanoma, treatment was initiated on day 7 after tumor inoculation (∼100 mm³) with nITC1, followed by nITC2 on day 8 (Figure 5A). A booster dose of nITC1 and a second nITC2 dose were administered on days 12 and 13, respectively, followed by nITC3 on day 18. Tumor growth was monitored until day 24. Control groups included individual nITC components, nITC(1+2), and a standard vaccine (TRP2/gp100 + CpG). Compared with monotherapies and the standard vaccine, both nITC(1+2) and nITC(1+2+3) significantly suppressed tumor growth, with the triple combination showing the greatest and most sustained efficacy (Figure 5B). Flow cytometric analysis revealed markedly increased intratumoral CD8^+^ and CD4^+^ T-cell infiltration in the nITC(1+2+3) group (Figure 5C), accompanied by significant reductions in PMN and M-MDSCs (Figure 5D), consistent with effective MDSC depletion by nITC3. In addition, nITC(1+2+3) and nITC3 increased the M1/M2 macrophage ratio in tumors and lymph nodes (Figure 5E; Supplementary Figure 5(I)), indicating polarization toward a proinflammatory phenotype. The triple therapy significantly increased CD44⁺CD62L⁺ central memory CD4^+^ and CD8^+^ T-cells (Figure 5F) and enhanced antigen-specific IFN-γ secretion following *ex vivo* restimulation with TRP2 and gp100 peptides (Figure 5G). In a single sequential treatment study (Figure 5H), nITC(1+2+3) provided superior tumor control (Figure 5I) and extended survival beyond 43 days, compared with ≤36 days for nITC1 or nITC(1+2) (Figure 5J). An increased number of tumor-infiltrating CD4^+^ and CD8^+^ T-cells (Figure 5K) was observed in the combination treatment. Tetramer staining for TRP2 and gp100 specific CD8^+^ T-cells revealed a significant expansion of tumor-specific T-cells in the triple therapy group (Figure 5L), correlating with improved tumor suppression and survival. Also, an increase in NK cells (Figure 5M) and their CD69 expression (Supplementary Figure 5(II)A) was observed, suggesting that the combinatorial therapy not only enhances NK cell recruitment but also promotes their functional activation. The combination therapy enriched central and effector memory, and naïve CD4^+^ and CD8^+^ T-cell subsets within tumors (Supplementary figure 5(II)C-D). To evaluate the safety of nITC(1+2+3), H&E staining of major organs (liver, kidney, spleen, heart, lung, lymph node, and brain) was performed 21 days after treatment. No observable pathological abnormalities were detected, indicating the absence of treatment-related toxicity (Supplementary Figure 5(III)).

**Figure 5.**
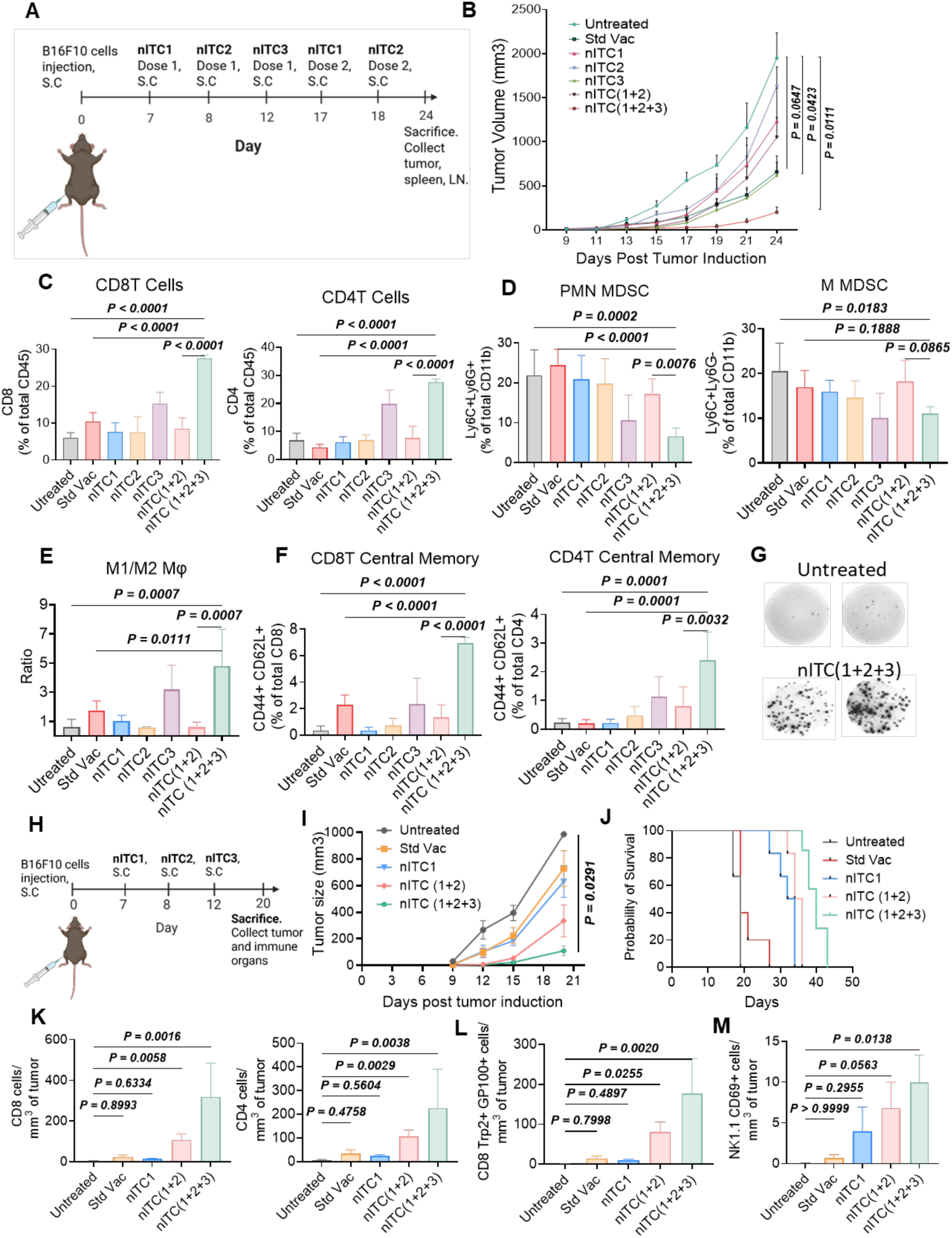
Combinatorial activation of nITC1, nITC2, and nITC3 enhances anti-tumor immunity in a murine melanoma model. A) Experimental design showing the treatment schedule for multiple dosing of nano-Immunotherapeutics (nITC1, nITC2, nITC3) in C57BL/6J mice bearing subcutaneous B16F10 melanoma tumors where nITC1 is n-CMZ(1:4:1), n-C=0.5 mM, n-M= 2 mM and n-Z= 0.5 mM; nITC2: hydrogel loaded with 10 ng of IL-15; nITC3: hydrogel loaded with 100 mg/kg 5-FU. B) Tumor growth curves of tumor-bearing animals treated as described; n=5; Experimental units=2. C) Bar graph showing frequency of intratumoral CD8⁺ T cells (left) and CD4⁺ T cells (right) on day 24. D) Bar graphs showing frequency of intratumoral PMN-MDSCs (left) and M-MDSCs (d, right). E) M1/M2 ratio, and central memory CD8⁺ (F, left) and CD4⁺ T cells (F, right), G) Representative ELISpot images of IFN-γ secreting splenocytes from untreated and nITC (1+2+3) treated mice following TRP2 restimulation. H) Schematic depicting the single-dose treatment schedule. I) Tumor volume measurements in tumor-bearing animals treated as described; n=5; Experimental units=2. J) Survival analysis of tumor-bearing animals treated as described. n=5; Experimental units=2. K) Bar graphs showing number of CD8^+^ (left) and CD4^+^ T cell (right) infiltrated into the tumor. L) Bar graphs showing the number of CD8^+^ Trp2^+^ gp100^+^ cells in tumor. M) Bar graphs showing the number of NK cells infiltrated into the tumor. Data are represented as mean ± SEM. P values were generated using ordinary one-way ANOVA with Tukey’s multiple comparison’s test (B, C, D) and Dunn’s multiple comparison’s test (E, F, K, L, M) for comparison. n = number of animals per group. Panel A and H created using BioRender.

### Combinatorial nITC treatment inhibits lung metastasis in mice

Lastly, we evaluated the therapeutic potential of the nITC(1+2+3) platform in a metastatic melanoma model, where MDSCs play a critical role^47,48^. Melanoma lung metastases were established by intravenous injection of melanoma cells (5 × 10⁵ cells), followed by treatment with nITC(1+2+3) according to the schedule shown in Figure 6A. Lungs were harvested on day 16 for analysis. Gross examination revealed extensive melanoma micrometastases in the lungs of untreated mice, indicating widespread tumor dissemination (Figure 6B, top panel). Lungs from nITC(1+2+3) treated animals exhibited a striking significant reduction in metastatic foci, with large areas of lung parenchyma remaining intact (Figure 6B, bottom panel, Figure 6C). Histological analysis confirmed markedly reduced tumor infiltration in treated animals compared with controls (Figure 6D). Lung tissue showed significant depletion of both PMN and M-MDSCs following treatment, accompanied by a marked increase in activated CD8⁺ CD69⁺ T-cells (Figure 6E-F). There was a significant reduction in circulating PMN-MDSCs (Figure 6G) and M2 macrophages (Figure 6H), while increasing total CD3⁺ T-cells, including both CD8⁺ and CD4⁺ subsets (Figure 6I-J). Treated mice exhibited increased frequencies of CD62L⁺CD44⁺ memory T-cells (Figure 6K) and reduced CD4⁺FoxP3⁺ regulatory T-cells (Figure 6L). These findings demonstrate that nITC(1+2+3) effectively suppresses metastatic progression by depleting MDSCs, reducing immunosuppressive macrophages and Tregs, and promoting robust cytotoxic and memory T-cell responses both locally and systemically. These results highlight the therapeutic potential of multi-component nanoparticle immunotherapy for controlling metastatic disease.

**Figure 6.**
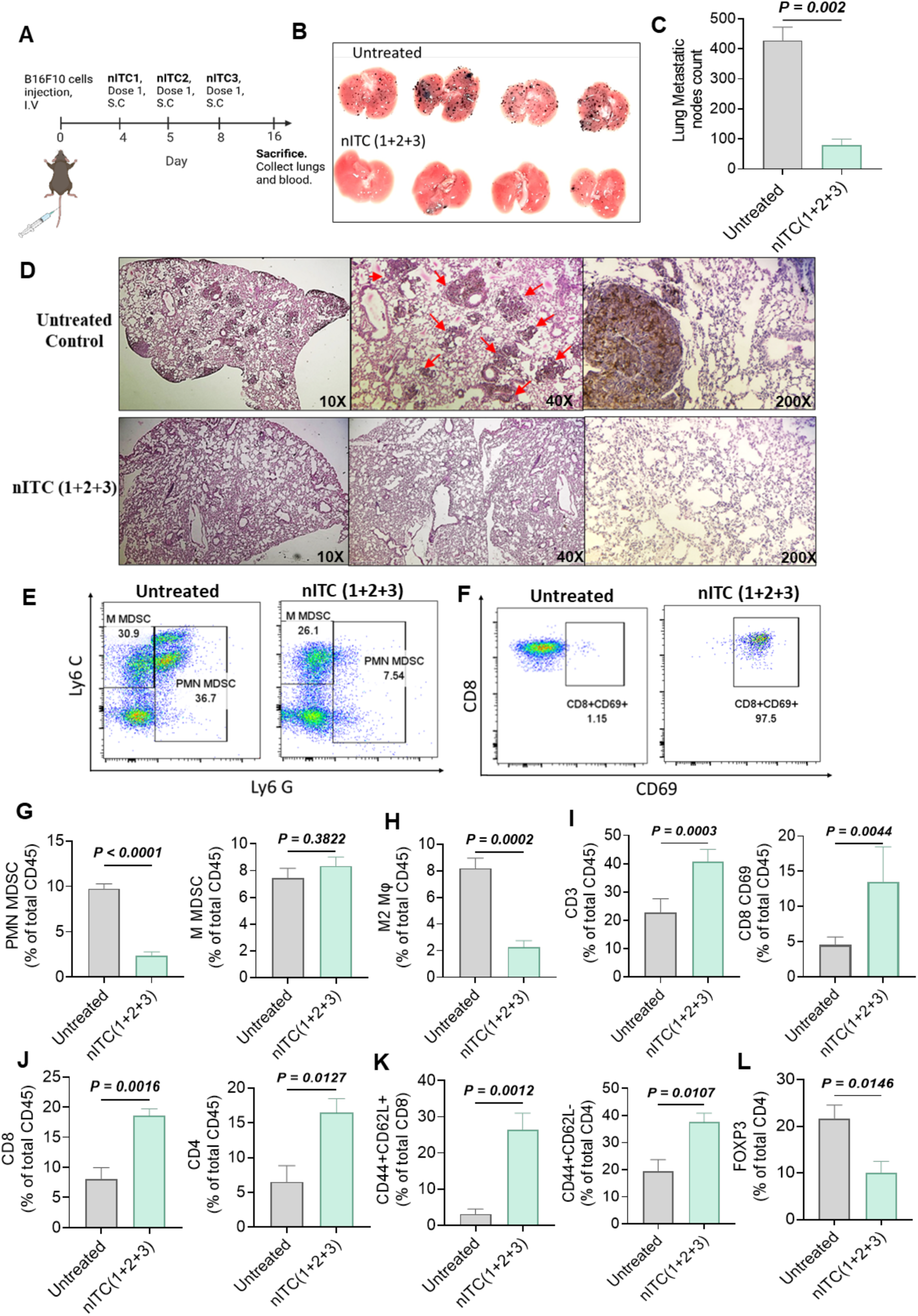
nITC combination inhibited lung metastasis in mice. A) Schematic representation of the experimental timeline. B) Representative images of lungs isolated from untreated controls and nITC-treated mice; n=4; Experimental units=2 C) Bar graph representation of lung metastatic nodules in untreated v/s nITC-treated mice. D) Histological analysis of lung sections (H&E staining) comparing control (top) and treated (bottom) groups are shown at multiple magnifications (10X, 40X and 200X). Dense tumor infiltration is highlighted by red arrows. E) Representative flow cytometry plots showing the frequencies of M-MDSCs (CD11b⁺Ly6C⁺Ly6G⁻) and PMN-MDSCs (CD11b⁺Ly6C⁺/− Ly6G⁺) in lungs from untreated (left) and nITC(1+2+3) treated (right) mice. F) Representative flow cytometry plots of activated CD8⁺ T cells (CD8⁺CD69⁺) in lungs from untreated and nITC(1+2+3) treated mice. G) Bar graph showing percentage expression of PMN-MDSCs and M-MDSCs, (H) M2 macrophages, (I) CD3⁺ T cells in blood. J) Bar graph showing frequency of circulating lymphocyte subsets. (K) Bar graph showing frequency of circulating CD4⁺ central memory T cells (left) and CD8⁺ central memory T cells (right). (L) Bar graph showing frequency of CD4⁺FoxP3⁺ regulatory T cells. Data are represented as mean ± SEM. Statistical analysis was performed using unpaired two-tailed t-test. n = number of animals per group. Panel a created using BioRender.

## Discussion

This study demonstrates that effective and durable anti-tumor immunity requires a rationally engineered trimodal nano-immunotherapy integrating three complementary strategies: enhanced immune cell priming (nITC1), sustained IL-15 release (nITC2), and TME remodelling (nITC3). Collectively, these modalities reprogram the anti-tumor immune response, resulting in more robust and durable immunity than conventional single-agent vaccination. A key innovation of this study is the rational engineering of a tri-metal phosphate nano-adjuvant that leverages the complementary immunological functions of Ca^2+^, Zn^2+^, and Mn^2+^ ions. Mn²⁺ is a potent activator of the cytosolic cGAS-STING pathway, driving type-I interferon and enhancing DC antigen presentation. Consistent with this, we show that Mn-containing nanoparticles markedly upregulated IFN-β and CD80/CD86 expression in DCs, underscoring their role in innate immune sensing and co-stimulation. Zn²⁺, known to promote Th1 differentiation and IFN-γ secretion, was shown here to significantly increase IFN-γ release from human PBMCs, thereby potentiating adaptive effector responses. In parallel, Ca²⁺, a central mediator of T-cell receptor signaling facilitated robust antigen-specific proliferation of CD8^+^ T-cells. By combining these three ions in a single hydrogen phosphate nanoparticle (nCMZ), we derived a unique multifunctional micronutrient nano-adjuvant for cancer vaccination. A complementary requirement for effective cancer vaccination is the persistence of cytokine signaling. IL-15 occupies a unique niche in immunotherapy due to its capacity to expand and sustain memory-like phenotypes of innate and adaptive cytotoxic lymphocytes, without driving regulatory T-cells. However, systemic administration of free IL-15 is limited by its short half-life (< 60 mins), high peak serum concentrations, and systemic toxicity. Recent clinical studies have demonstrated that recombinant IL-15 can promote NK- and T-cell activation when combined with immune checkpoint blockade^49^. Our data show that nITC2 provided sustained cytokine release, transiently enhanced NK and CD8⁺ T-cell activation, and prolonged systemic immune stimulation. These findings support nITC2 as a safe and effective cytokine delivery platform for cancer immunotherapy. Another key challenge we addressed was immune suppression in the TME. High MDSC burden correlates with poor prognosis and therapeutic resistance across multiple cancer. Sustained release of 5-FU from nITC3 effectively reduced frequencies of both PMN- and M-MDSCs, lowered TGF-β expression, and increased M1 macrophage polarization.

One of the key aspects of our work is the translational benefit of integrating all three modules using nanotechnology. The combined system of nITC (1+2+3) produced the most profound effect in tumor reduction and reduced pulmonary metastases. From a translational perspective, this trimodal strategy offers several advantages: a micronutrient based nanoadjuvant that activates both innate and adaptive immunity, a biocompatible hydrogel for sustained IL-15 release to support durable effector and memory responses, and TME remodeling through myeloid cell depletion. Together, these findings provide a blueprint for developing integrated nano-immunotherapies adaptable to diverse tumor types and personalized antigen repertoires. Consistent with the translational potential of biomaterial-based immunotherapies, the recent first in human WDVAX scaffold vaccine trial demonstrated the clinical feasibility of sustained local immune modulation using a microporous scaffold loaded with GM-CSF, CpG, and autologous tumor lysate^50^. Our findings further extend this concept by integrating antigen priming, cytokine support, and TME remodelling using a cost-effective immunotherapy.

Nonetheless, limitations of our study include reliance on a single preclinical melanoma model, lack of long-term safety and biodistribution data, and the need to test synergy with checkpoint blockade in heterogeneous tumor settings. Future studies should evaluate these aspects, including pharmacokinetics, biodistribution, toxicity, and biomarker-driven stratification of tumor types.

## Declarations

### Ethics approval

Animal procedures were approved either by the Institutional Animal Ethics Committee of Amrita Institute of Medical Sciences, Kochi, India (Approval No. IAEC/2021/1/4) or by the Cantonal Veterinary Authorities of Geneva and Vaud Cantons (Authorisation No. VD3496 and VD3772). Buffy coats from healthy human blood donors were collected following approval from the Institutional Ethics Committee (IEC), Amrita Institute of Medical Sciences, Kochi, India (Approval No. IAEC-AIMS-2020-NANO-162).

### Consent for publication

Not applicable. This manuscript does not contain any individual person’s data or identifiable images. All authors have read the final version of the manuscript and give their consent for publication.

### Availability of data and material

The data analyzed and presented in this study are available upon request to the corresponding author.

### Authors’ Contributions

Planning and supervising: PR, CJ and MK. Experimenting: ZTZ, SNM, MS, AKS, AGC, HEA, SR, AG, AMN and SN. Writing: ZTZ, AGC, HEA, PR, CJ and MK.

## Acknowledgments

The authors thank the Central Laboratory Animal Facility, AIMS, Kochi, for conducting the *in vivo* experiments, and the Amrita School of Nanosciences and Molecular Medicine (ASNSMM), Amrita Vishwa Vidyapeetham, for providing the necessary infrastructure and institutional support. The authors thank the University of Lausanne animal and flow cytometry facilities for excellent assistance.

