## Supplementary Figures for "Integrated nano-immunotherapy for synergizing cancer vaccination with sustained cytokine release and tumor microenvironment modulation"

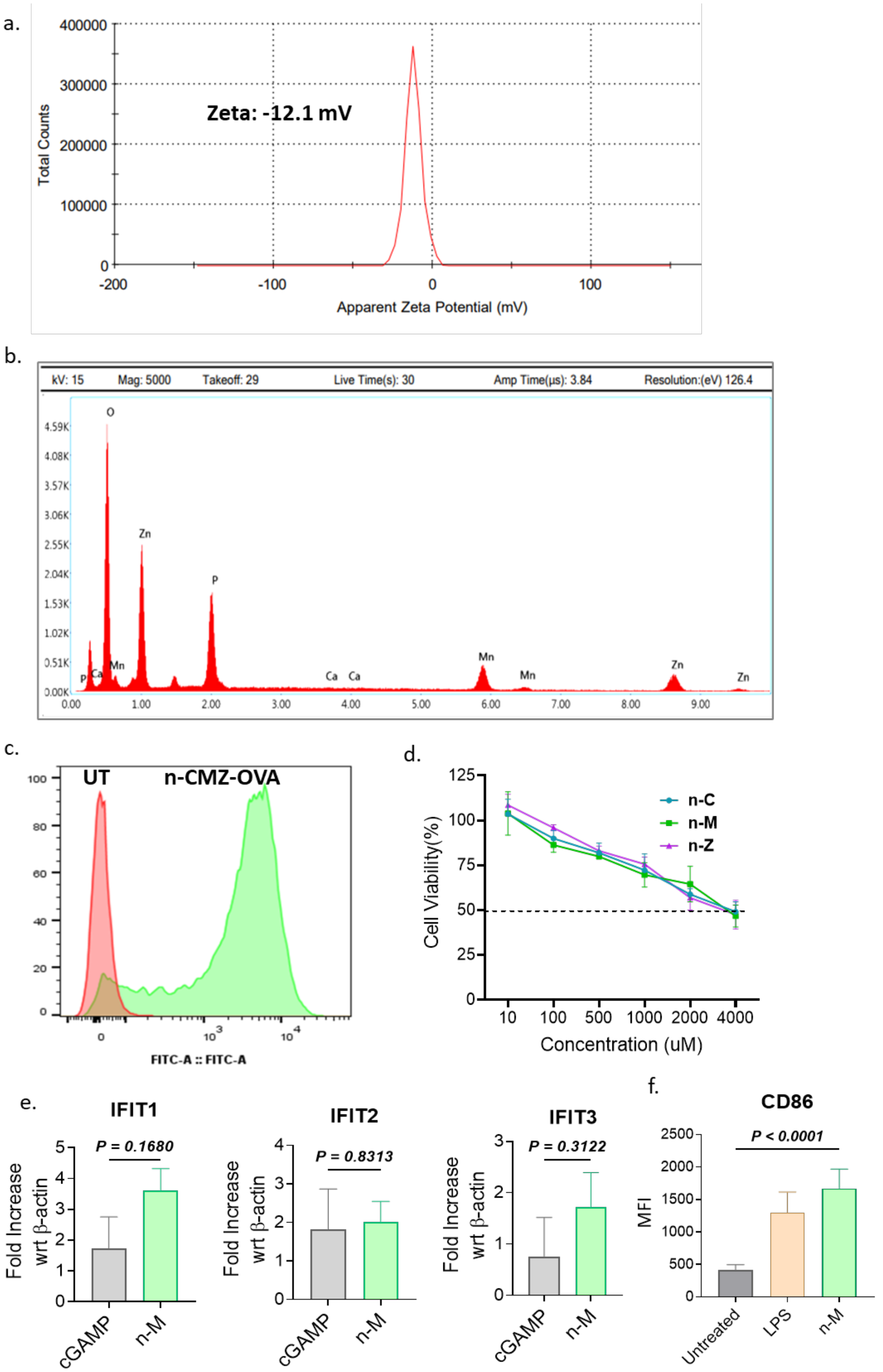


**Supplementary Figure 1. CMZ Characterization and immunological evaluation**. a) Zeta potential measurement of nanoparticles reveals a negative surface charge of -31.8 mV. b) Energy dispersive X-ray spectroscopy (EDS) confirms the elemental composition of the nanoparticles, showing peaks for Zn, O, P, Ca, and Mn, consistent with a mixed metal phosphate formulation. c) Flowcytometric confirmation of DC uptake of FITC-OVA conjugated nITC1 nanoparticles. d) Dose-dependent cytotoxicity of nanoparticles containing individual elements in the mouse PBMCs e) Expression analysis of IFIT1, IFIT2, and IFIT3 genes in DCs treated with cGAMP and 100 μM of n-M. f) Bar graph showing increase in activation of CD86 in DCs treated with 100 μM n-M compared to untreated and LPS by flow cytometry Data are represented as mean ± SEM. P value in (e) was performed using unpaired two-tailed t-test. P values of (f) were generated using ordinary one-way ANOVA with Dunnet's multiple comparison's test for comparison.

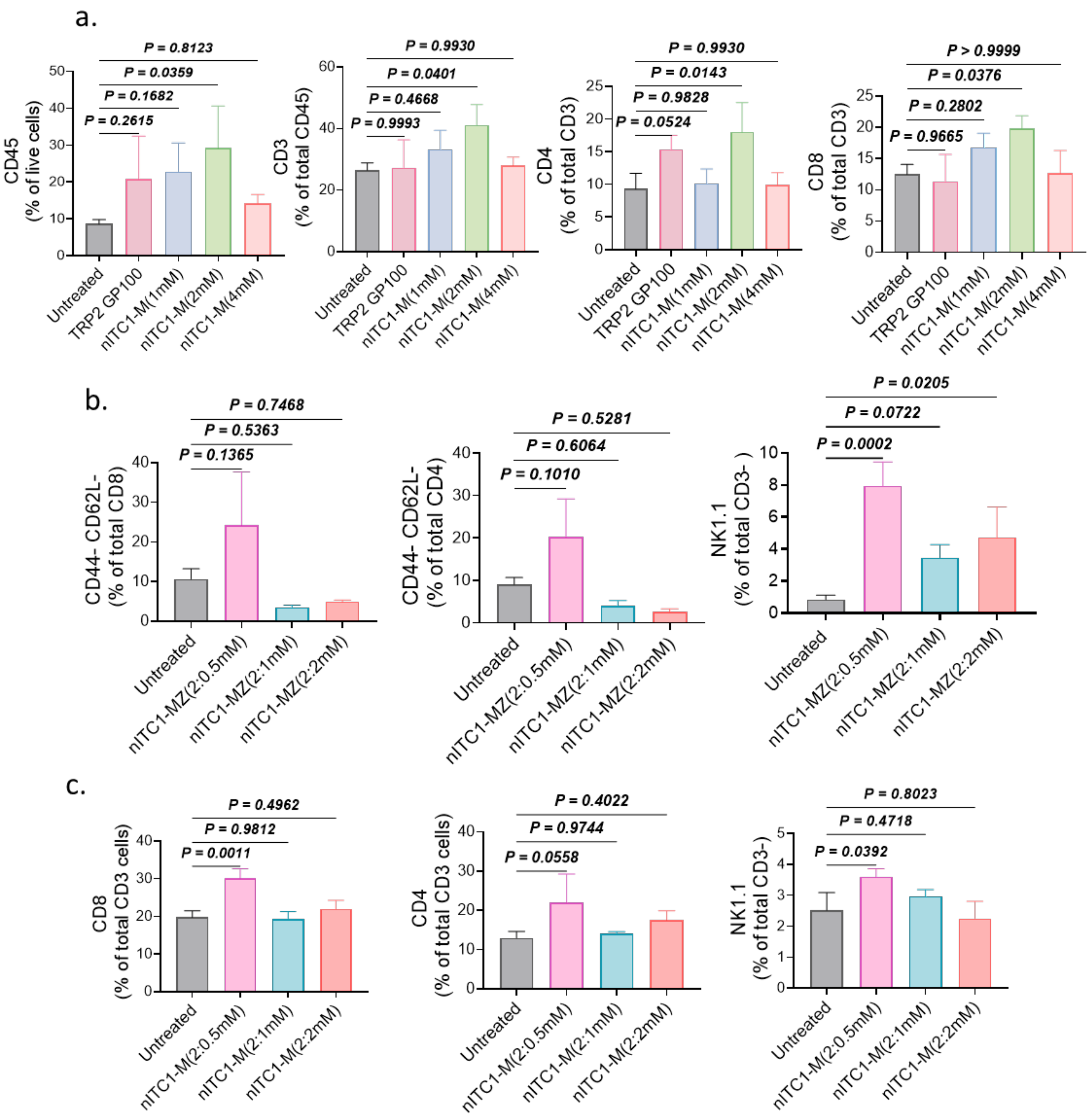


**Supplementary Figure 2(i). Immune cell profiling in tumor and spleen following nanoparticle treatment.** a) Bar graphs showing frequency of CD45$^+$, CD3$^+$, CD4$^+$ and CD8$^+$ cells in tumor after 21 days in different nITC1-M groups. b) Bar graphs showing frequency of CD4$^+$, CD8$^+$ and NK cells in tumor after 21 days in in different nITC1-MZ groups. c) Bar graphs showing frequency of CD4$^+$, CD8$^+$ and NK cells in spleen after 21 days in different nITC1-MZ groups**.** Data are represented as mean ± SEM. P values were generated using ordinary one-way ANOVA with Dunnet’s multiple comparison’s test for comparison.

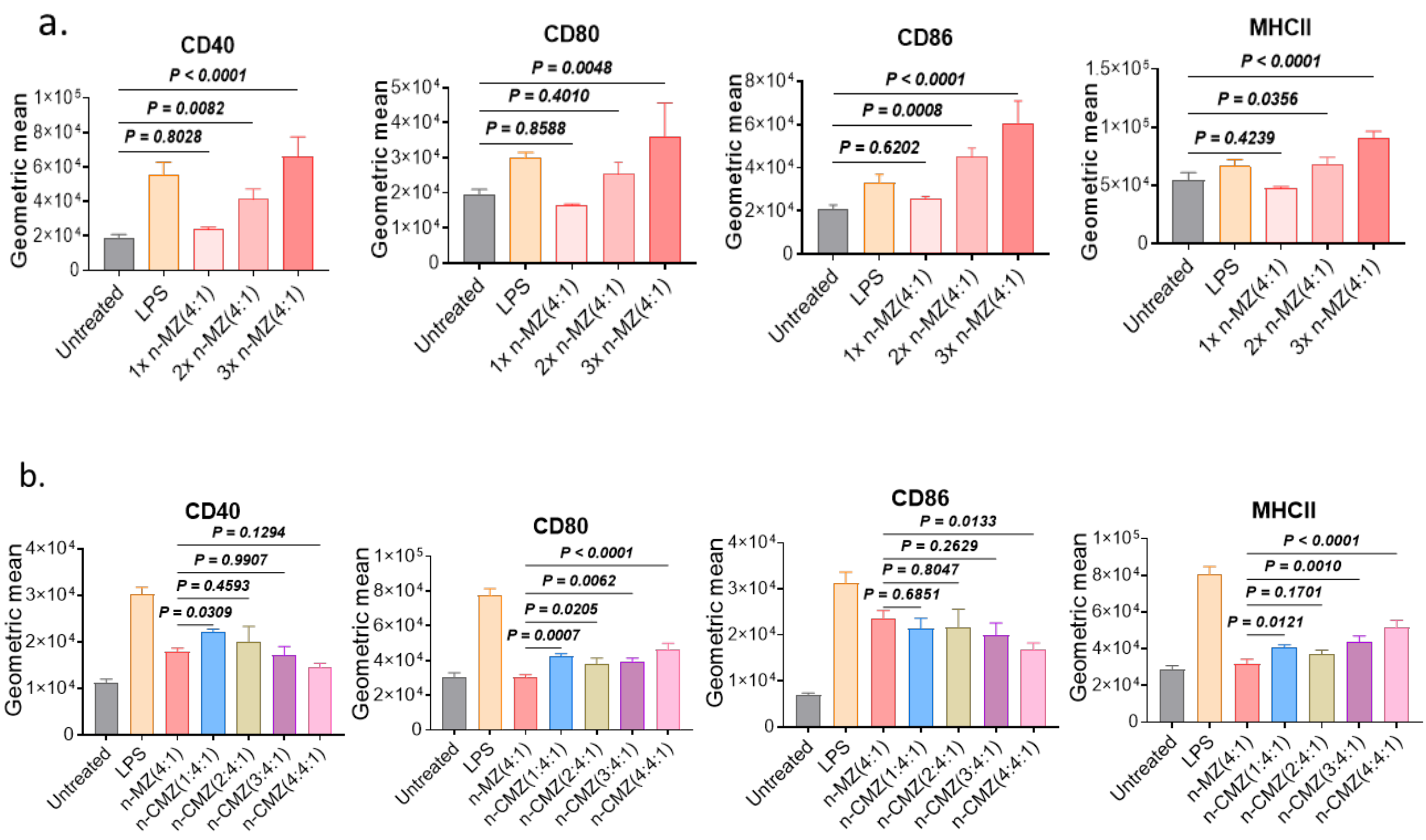


**Supplementary Figure 2(ii).** ***In vitro*** **activation of BMDCs following nanoparticle treatment.** a) Bar graph representation of CD40, CD80, CD86 and MHC-II in BMDCs treated *in vitro* with increasing dose of n-MZ Nanoparticles(4:1); X represents increasing dose (µM), 1X n-MZ(4:1) contains 25 µM n-M and 6.25 µM n-Z, 2X n-MZ(4:1) contains 50 µM n-M and 12.5 µM n-Z, 3X n-MZ(4:1) contains 100 µM n-M and 25 µM n-Z. b) Bar graph representation of CD40, CD80 , CD86 and MHC-II in BMDCs *in vitro* treated with n-MZ(4:1) in combination with different concentration of n-C, where n-CMZ (1:4:1) contains 25 µM n-C, 100 µM n-M and 25 µM n-Z; n-CMZ (2:4:1) contains 50 µM n-C, 100 µM n-M and 25 µM n-Z; n-CMZ (3:4:1) contains 75 µM n-C, 100 µM n-M and 25 µM n-Z; n-CMZ (4:4:1) contains 100 µM n-C, 100 µM n-M and 25 µM n-Z. Data are represented as mean ± SEM. P values were generated using ordinary one-way ANOVA with Dunnet's multiple comparison's test for comparison.

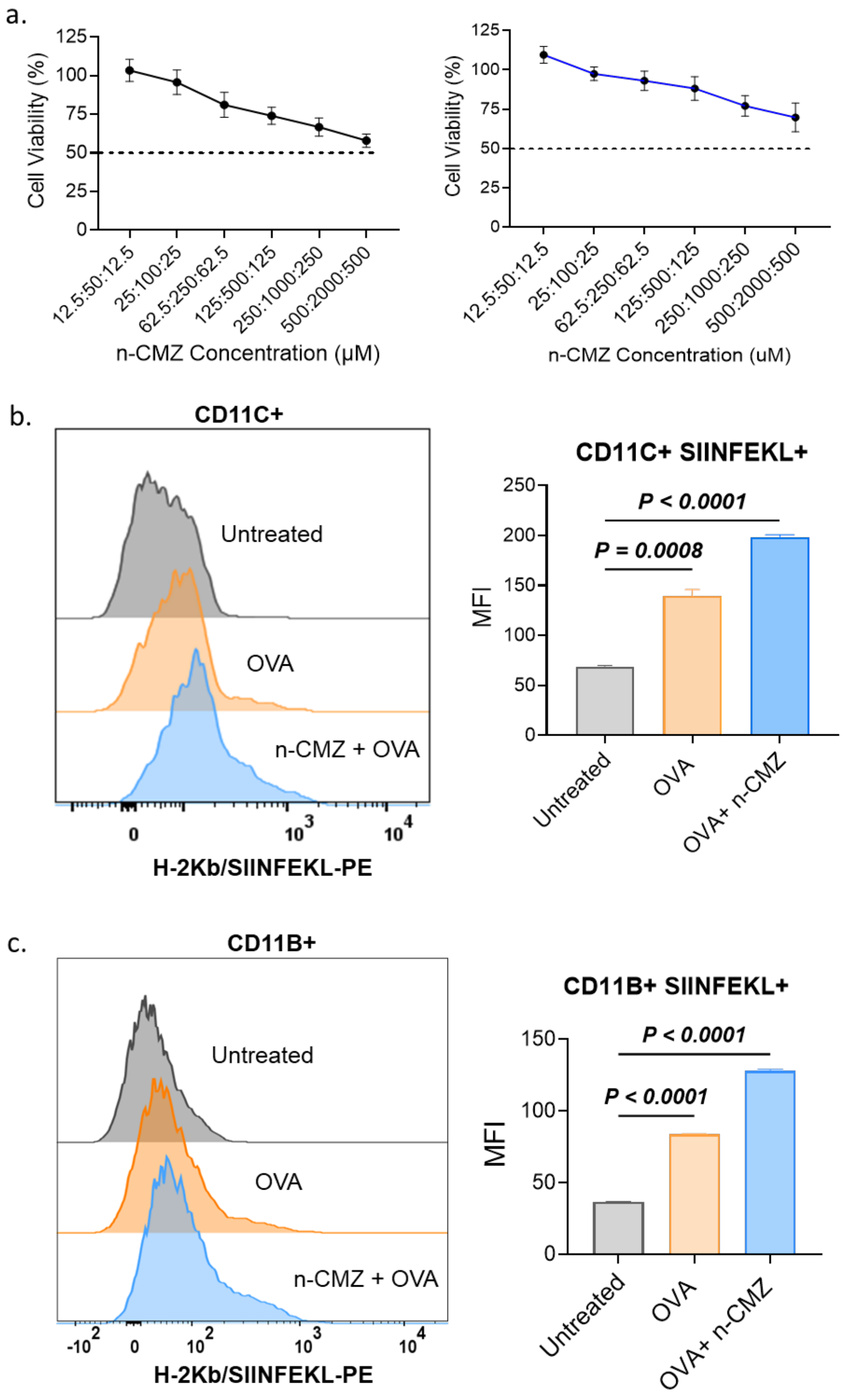


**Supplementary Figure 2(iii)**. **Cytotoxicity and cellular uptake of n-CMZ nanoparticles**. a) Cytotoxicity of n-CMZ (1:4:1) with increasing concentration of different elements in human PBMCs (left) and mouse DCs (right): C=12.5,25,62.5,125,250,500 µM; M=50,100,250,500,1000,2000 µM; Z=12.5,25,62.5,125, 250,500 µM respectively. b) Overlayed histograms depict OVA-PE fluorescence intensity in CD11c$^{+}$ DCs either untreated, treated with OVA alone, or with OVA+CMZ nanoparticles (left); corresponding mean fluorescent intensity is shown as bar graph (right). c) Overlayed histograms depict OVA-PE fluorescence intensity in CD11b$^{+}$ myeloid cells with the same treatment groups (left)**.** corresponding mean fluorescent intensity is shown as bar graph (right). Data are represented as mean ± SEM. P values was generated using ordinary one-way ANOVA with Tukeys's multiple comparison's test for comparison.

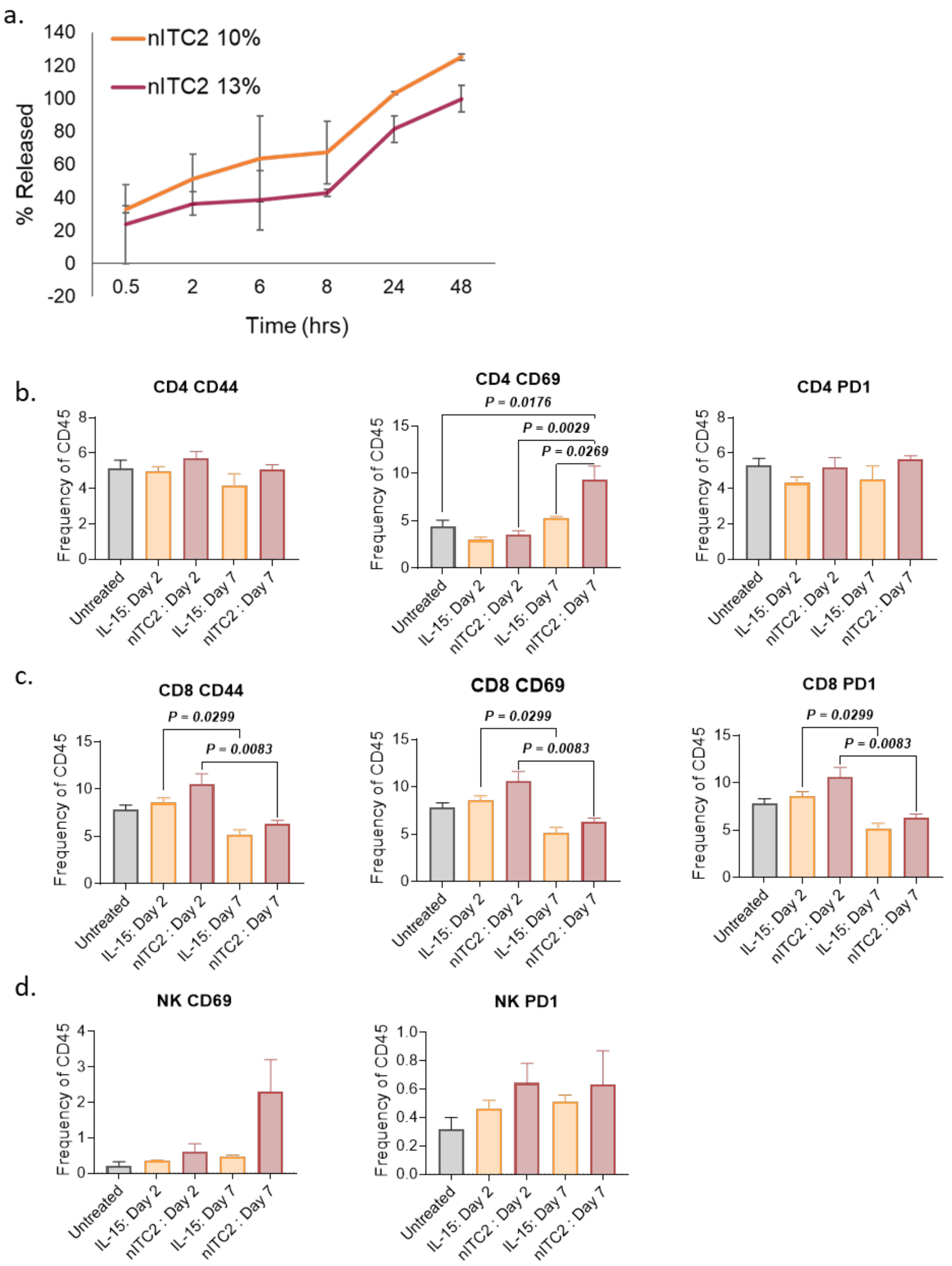


**Supplementary Figure 3(i). nITC2-mediated protein release and activation of T cell and NK cell responses.** a) *In vitro* release profile showing the percentage release kinetics of BSA from nITC2 of two different gel percentage loaded with 500 µg of model protein BSA. b-c) Bar graphs showing the frequency of $CD4^{+}$, $CD8^{+}$ T cell and NK cell activation markers (CD69, CD44, PD1) on day 2 and day 7 post-treatment in lymph nodes. Data are represented as mean ± SEM. P values were generated using one way ANOVA with Tukey's multiple comparison's test for comparison.

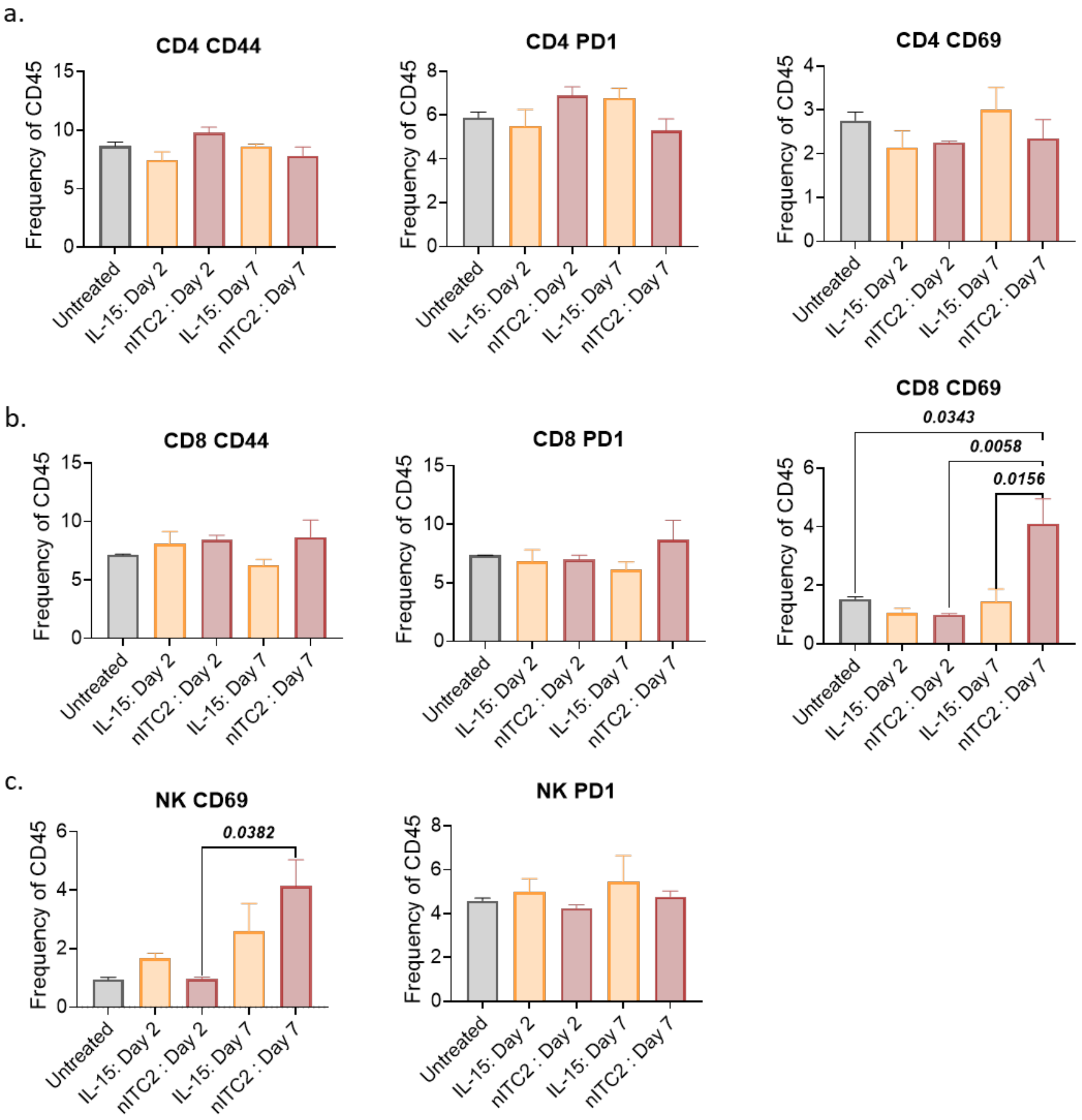


**Supplementary Figure 3(ii). nITC2-mediated activation of T cell and NK cell responses in spleen.** a-c) Bar graphs showing the frequency of $CD4^+$, $CD8^+$ T cell and NK cell activation markers (CD69, CD44, PD1) on day 2 and day 7 post-treatment in mice spleen. Data are represented as mean ± SEM. P values was generated using one way ANOVA with Tukey's multiple comparison's test for comparison.

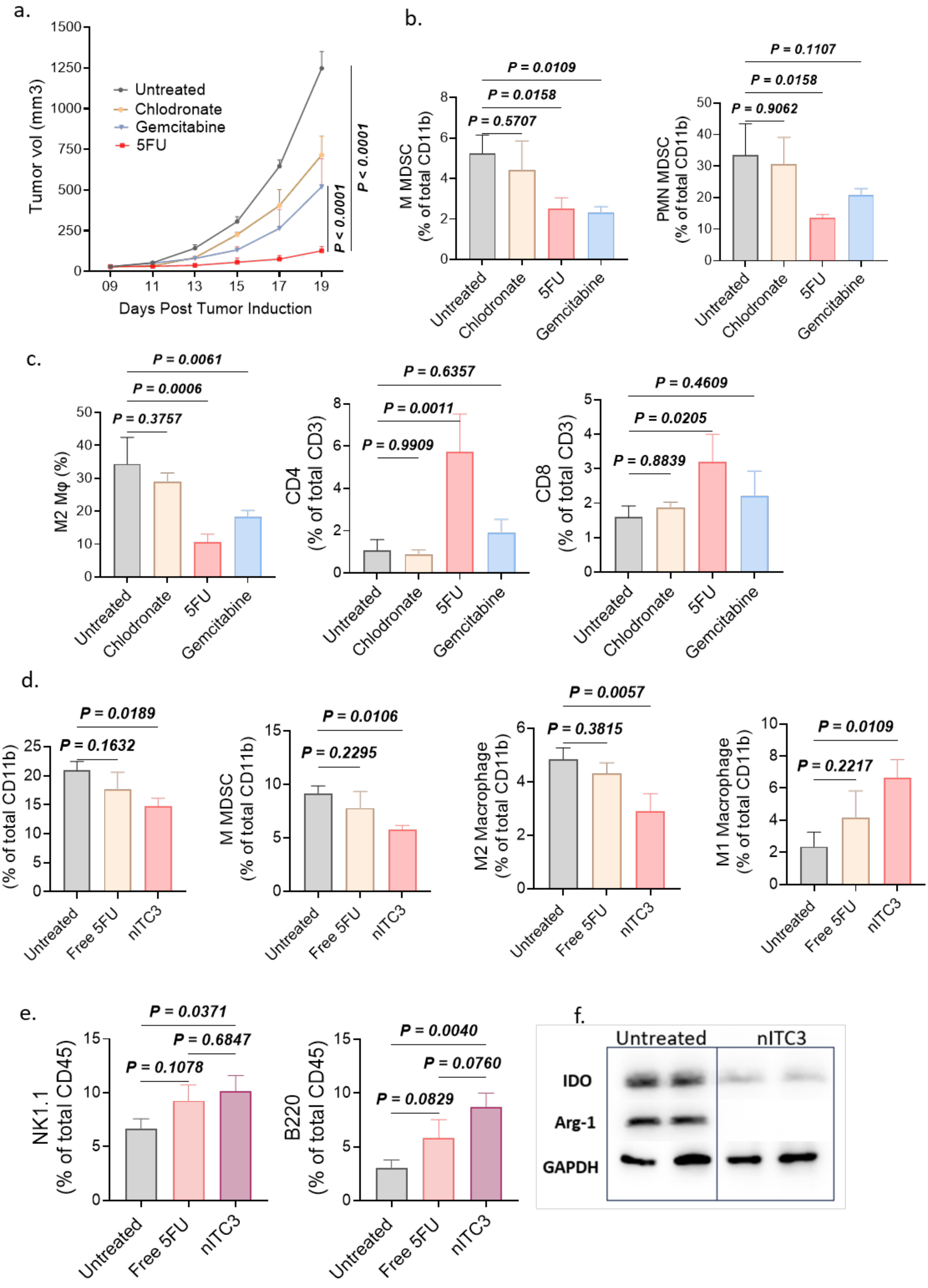
a.
Tumor vol (mm3)
Untreated
Chlodronate
Gemcitabine
5FU
P < 0.0001
P < 0.0001
Days Post Tumor Induction
b.
M MDSC (% of total CD11b)
P = 0.0109
P = 0.0158
P = 0.5707
PMN MDSC (% of total CD11b)
P = 0.1107
P = 0.0158
P = 0.9062
Untreated
Chlodronate
5FU
Gemcitabine
c.
M2 Mφ (%)
P = 0.0061
P = 0.0006
P = 0.3757
CD4 (% of total CD3)
P = 0.6357
P = 0.0011
P = 0.9909
CD8 (% of total CD3)
P = 0.4609
P = 0.0205
P = 0.8839
d.
PMN MDSC (% of total CD11b)
P = 0.0189
P = 0.1632
M MDSC (% of total CD11b)
P = 0.0106
P = 0.2295
M2 Macrophage (% of total CD11b)
P = 0.0057
P = 0.3815
M1 Macrophage (% of total CD11b)
P = 0.0109
P = 0.2217
Untreated
Free 5FU
nITC3
e.
NK1.1 (% of total CD45)
P = 0.0371
P = 0.6847
P = 0.1078
B220 (% of total CD45)
P = 0.0040
P = 0.0760
P = 0.0829
f.
Untreated
nITC3
IDO
Arg-1
GAPDH

**Supplementary Figure 4. Antitumor efficacy and immune modulation by nITC3 v/s free 5-FU treatment.** a) Tumor growth curves comparing untreated, 5-FU, gemcitabine, chlodronate treated animals (n=3), showing enhanced tumor suppression with 5-FU (left). Bar graphs showing the frequency of PMN-MDSC and M-MDSC in the tumor. b) Bar graph showing the frequency of M2 macrophages, $CD4^+$ and $CD8^+$ T cell populations in the tumor. c) Bar graphs showing the frequency of PMN-MDSC, M-MDSC, M2 macrophages and M1 macrophages in spleen of free 5-FU v/s nITC3 treated animals. d) Bar graphs representation of frequency of $NK1.1^+$ and $B220^+$ cells in the tumor. e)Western blot analysis showing reduced expression of immunosuppressive enzymes IDO and Arg-1 in nITC3-treated tumors compared to untreated controls. GAPDH was used as control. Data are represented as mean ± SEM. P values in (c) was generated using two-way ANOVA with Tukey's multiple comparison test. P values in (b-e) were generated using ordinary one-way ANOVA with Dunnet's multiple comparison's test for comparison.

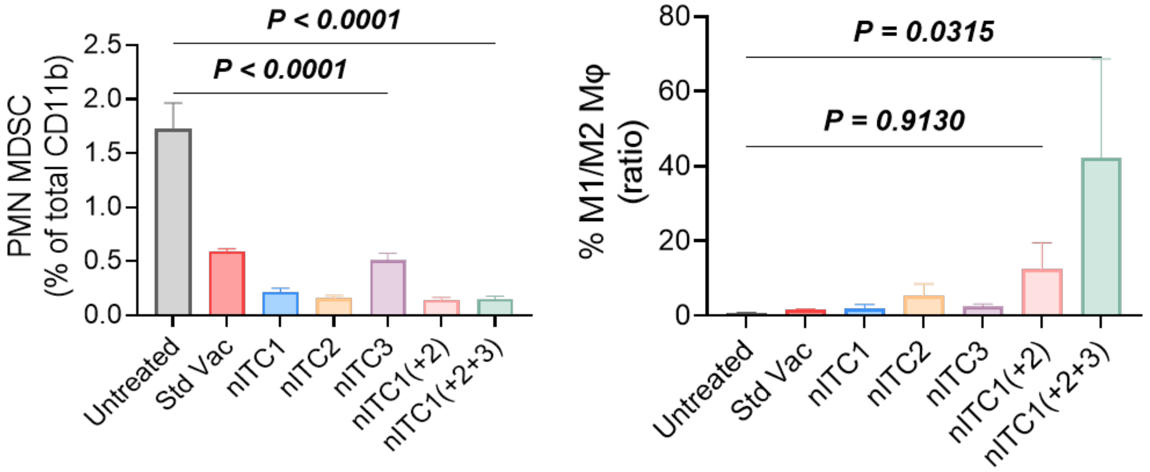


**Supplementary Figure 5(i)**. **Myeloid cell composition in lymph nodes.** Bar graphs showing the frequency of PMN-MDSC (left) and ratio of M1/M2 macrophages (right) in lymph nodes. Data are represented as mean ± SEM. P values were generated using ordinary one-way ANOVA with Dunnet's multiple comparison's test for comparison.

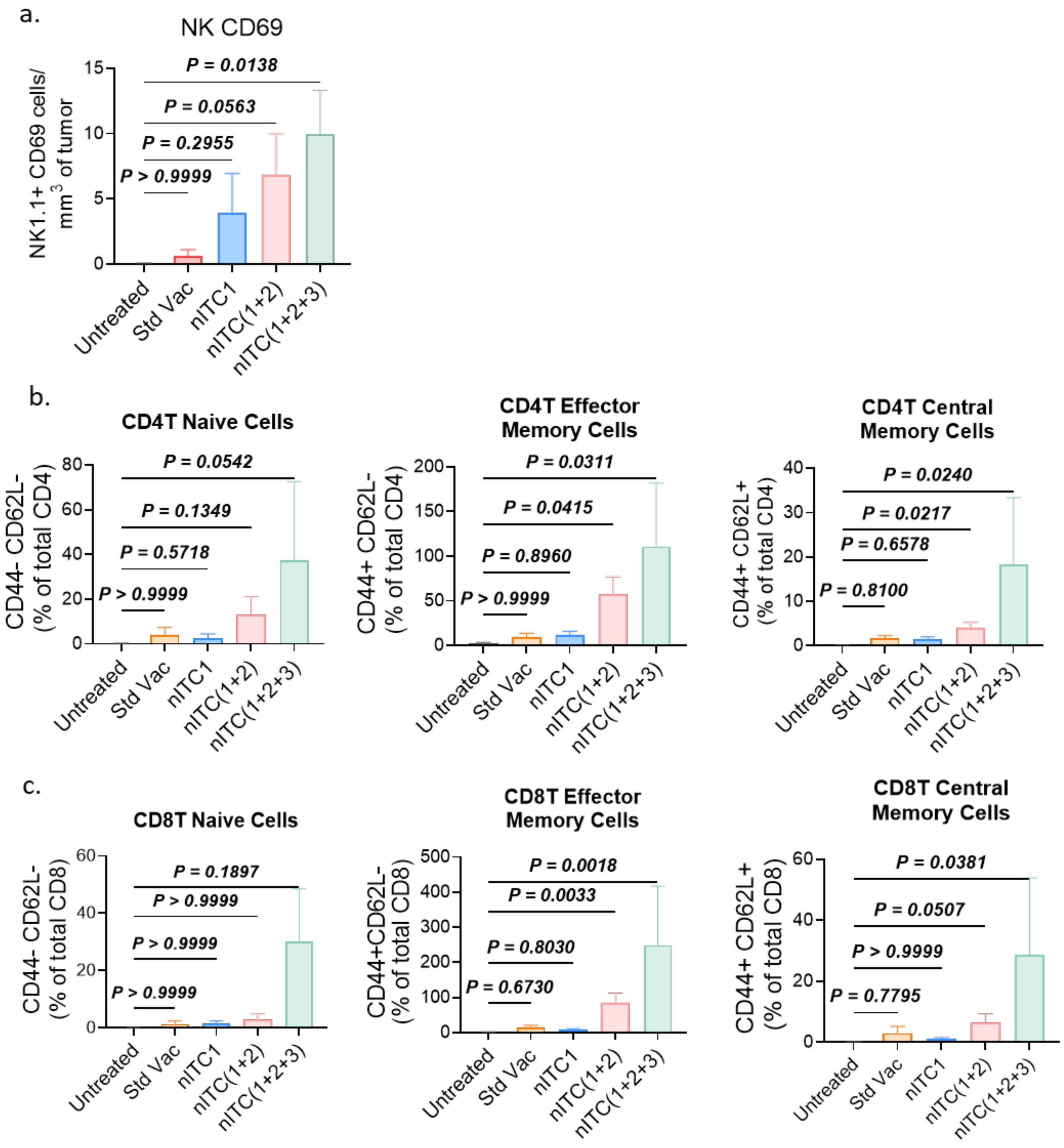


**Supplementary Figure 5(ii). Immune cell activation and T cell subset distribution following combination treatment** a) Bar graph showing number of activated NK cells (NK1.1$^+$ CD69$^+$) infiltrated into the tumor with combination treatment. b) Bar graphs showing frequencies of naïve, effector memory, and central memory CD4$^+$ T cells. c) Bar graphs showing frequencies of naïve, effector memory, and central memory CD8$^+$ T cells. Data are represented as mean ± SEM. P values were generated using ordinary one-way ANOVA with Kruskal Wallis multiple comparison's test.

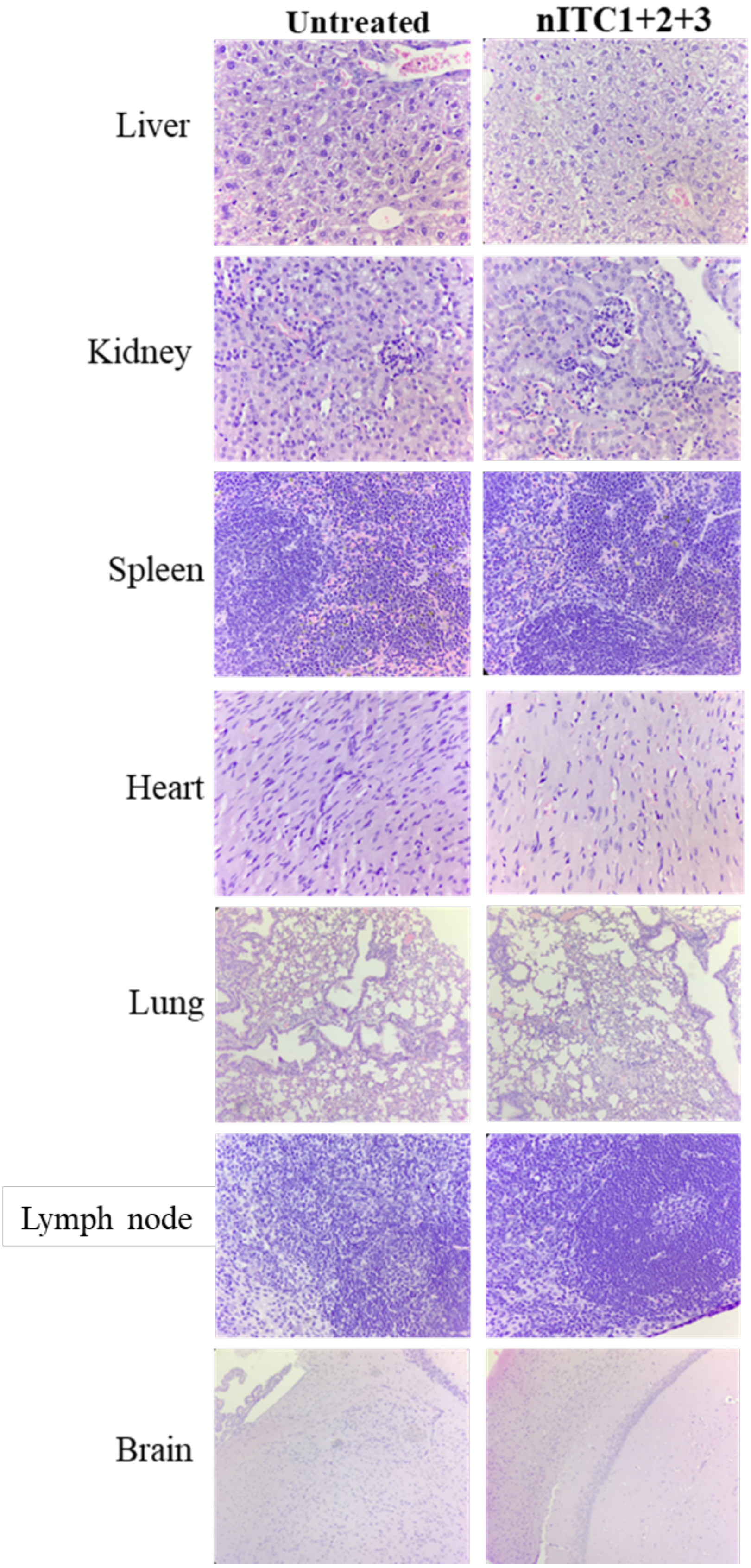


**Supplementary Fig 5(iii)**. Haematoxylin and eosin staining images of main organs (liver, kidney, spleen, heart, lung, lymph node and brain) of C57BL/6J mice after 21 days of administration of nITC (1+2+3) as compared to untreated controls. Scale bar, 60 µm.
